# CRISPR-Induced NUT Suppression Promotes Differentiation and Enhances Trop2-Targeted Immunocytokine Response in NUT Carcinoma

**DOI:** 10.1101/2025.10.28.685224

**Authors:** Juyoung Choi, Daesik Kim, Junghan Oh, Hee Geon Park, Hwa Lin Yi, Yurimi Lee, Hoyeon Jeong, Ji-Young Song, Minjung Sung, Eun Sol Chang, Kyungsoo Jung, Saehyung Lee, Young Kee Shin, Se-Hoon Lee, Christopher A French, Mi-Sook Lee, Yoon-La Choi

## Abstract

NUT carcinoma (NC) is an aggressive malignancy driven by *NUTM1* gene rearrangements with limited therapeutic options. Here, we show that direct suppression of *NUTM1* using CRISPR/Cas9 induces squamous-like differentiation and upregulates TROP2 expression in NC cells. Building on this finding, we developed a TROP2–interferon beta (IFN-β) mutein immunocytokine that selectively targets TROP2-expressing tumors. Combined *NUTM1* suppression and TROP2-targeted immunotherapy synergistically enhanced cytotoxic and immune-mediated responses *in vitro*. Transcriptomic and spatial analyses of NC patient tumors revealed that differentiation status correlates with TROP2 expression, upregulated immune pathways, and favorable clinical outcomes. Our results suggest that overcoming differentiation blockade not only alters tumor phenotype but also creates a more immune-permissive microenvironment. These findings highlight the therapeutic potential of sequential tumor reprogramming followed by targeted immunotherapy in treating NC and propose a broader strategy for overcoming differentiation blockade in fusion-driven cancers.

## Introduction

NUT carcinoma (NC), formerly known as NUT midline carcinoma, is a rare and aggressive malignancy characterized by chromosomal translocations in the NUT midline carcinoma family member 1 (*NUTM1*) gene.^1^ NC can affect individuals of all ages, from infancy to old age, with a nearly equal distribution between sexes.^2^ Despite advances in next-generation sequencing (NGS)-based diagnostics and increased awareness leading to higher detection rates, the prognosis of NC remains poor, with a median overall survival (OS) of only six to nine months. The poor prognosis is largely attributed to the frequent presence of metastases at diagnosis^3–6^ and the limited response to conventional chemotherapy and radiotherapy.^6,7^ Histologically, NC primarily presents as an undifferentiated or poorly differentiated carcinoma, typically composed of monomorphic primitive cells with hyperchromatic nuclei and scant cytoplasm with a sheet-like growth pattern without nuclear molding. Although poorly differentiated forms of NC are more prevalent, squamous differentiation or abrupt keratinization is observed in 33% of the patients.^3^ Squamous differentiation varies from subtle to prominent, with some patients showing clear keratinization.^8^ However, the clinical significance of intra- and inter-tumor histopathological heterogeneity in NC remains unclear.

Bromodomain-containing protein 4 (*BRD4*), a known *NUTM1* partner gene, is well characterized and plays a critical role in NC pathogenesis. Current NC the rapies largely rely on BET inhibitors (BETis) that target BRD4.^9,10^ However, clinical responses are modest and often accompanied by systemic toxicity due to nonselective BET protein targeting.^7,11–15^ To overcome these limitations, *NUTM1* itself has emerged as a more selective therapeutic target, given its restricted expression in post-meiotic spermatogenic cells.^16,17^ Direct *NUTM1* suppression using CRISPR/Cas9 gene editing offers improved specificity and reduced toxicity compared to BETis, thus presenting a promising strategy for the safer and more effective treatment of patients with NC.^18^

In the present study, *in vitro* CRISPR/Cas9-mediated *NUTM1* editing was found to inhibit tumor growth and induce the differentiation of NC tumor cells. This differentiation process significantly increased TROP2 expression, highlighting new opportunities for targeted therapy.^19,20^ Based on these findings, we classified the samples of NC patients into three categories according to their differentiation status and evaluated the relationship between differentiation, TROP2 expression, and clinical outcomes. This classification revealed an association between the differentiation status, prognosis, and presence of an immune-enriched microenvironment. To advance therapeutic strategies, we explored a sequential targeting strategy that integrates precise NUT suppression with TROP2-targeted immunocytokine therapy. This strategy not only enhances treatment specificity but also modulates the tumor microenvironment, thus fostering conditions favorable for additional therapeutic interventions.

## Results

### *NUTM1* gene editing suppresses *BRD4::NUTM1*-expressing cell proliferation and induces squamous-like differentiation

BRD4::NUT is an oncoprotein and a therapeutic target in NC. Inhibition of NUT expression using *NUTM1* siRNAs has been reported to halt tumor growth by inducing chromatin reprogramming, effectively reversing the oncogenic impact of the BRD4::NUT fusion protein through the downregulation of key oncogenes such as MYC.^21–23^ In this study, we explored whether targeting *NUTM1* using CRISPR/Cas9 could inhibit NC cell growth. Among the NC cell lines, RPMI2650 (nasal septum), HCC2429 (lung), JCM, and 10-15 were selected based on their relevance to the characteristic oncogenic features of NUT carcinoma. Detailed molecular profiles and tissue origins of these cell lines are summarized in Supplementary Table 1, while Supplementary Figure 1 illustrates their genomic alterations and fusion gene structures.

To investigate the functional impact of NUT suppression, we employed CRISPR/Cas9-mediated genome editing targeting multiple exons of the *NUTM1* gene, including the fusion-critical exon 3 and interaction associated exons 6 and 8. Among a panel of candidate sgRNAs, four (E3-2, E3-9, E6, and E8-1v2) demonstrated the highest editing efficiencies and were selected for subsequent experiments (Fig. 1A, Supplementary Fig. 2A-C, Supplementary Table 2). Cas9/sgRNAs ribonucleoprotein complexes were delivered using engineered virus-like particles (eVLPs), leading to a marked reduction in *NUTM1* mRNA and NUT protein expression levels by 50–60% within 72 hours post-transfection, relative to a control- C chemokine receptor type 5 (*CCR5*). This reduction in NUT expression was accompanied by decreased levels of oncogenic transcriptional targets, including TP63, MYC, and SOX2, particularly in HCC2429 and JCM1 cells (Fig. 1B). Functionally, CRISPR-mediated *NUTM1* disruption significantly impaired cell proliferation across all four NC cell lines tested, as demonstrated by WST assays (Fig. 1C). To further assess therapeutic potential under more physiologically relevant conditions, we employed a 3D spheroid culture model. After 13 days of eVLP-mediated delivery, spheroids edited with sgRNAs targeting E3-2, E3-9, and E6C (1:1 mixture of sg E6 and sgE8-1v2) exhibited significantly reduced growth compared to *CCR5* controls (p = 0.0242, p = 0.0002, and p < 0.0001, respectively) (Fig. 1D). Interestingly, NUT suppression not only inhibited tumor growth but also induced significant morphological changes in NC cells, including increased cytoplasmic volume and cell flattening and spreading (Fig. 1E and Supplementary Fig. 4C). To elucidate the underlying mechanisms of these morphological and proliferative alterations, we next examined markers of senescence, apoptosis, and differentiation. *NUTM1*-edited cells did not exhibit an increase in senescence-associated GFP expression compared to *CCR5* controls, indicating that senescence was not involved (Supplementary Fig. 3A). Apoptosis remained largely unchanged, although a slight increase in the pro-apoptotic fraction was observed in *NUTM1*-edited cells, indicating minor alterations in apoptotic signaling (Supplementary Fig. 3B-D). Notably, expression of genes associated with differentiation was upregulated following *NUTM1* editing (Supplementary Fig. 2D), supporting a differentiation-driven mechanism underlying the reduced proliferation. These findings suggest that CRISPR/Cas9-mediated *NUTM1* suppression primarily promotes differentiation rather than inducing senescence or apoptosis in NC cells.

**Figure 1.**
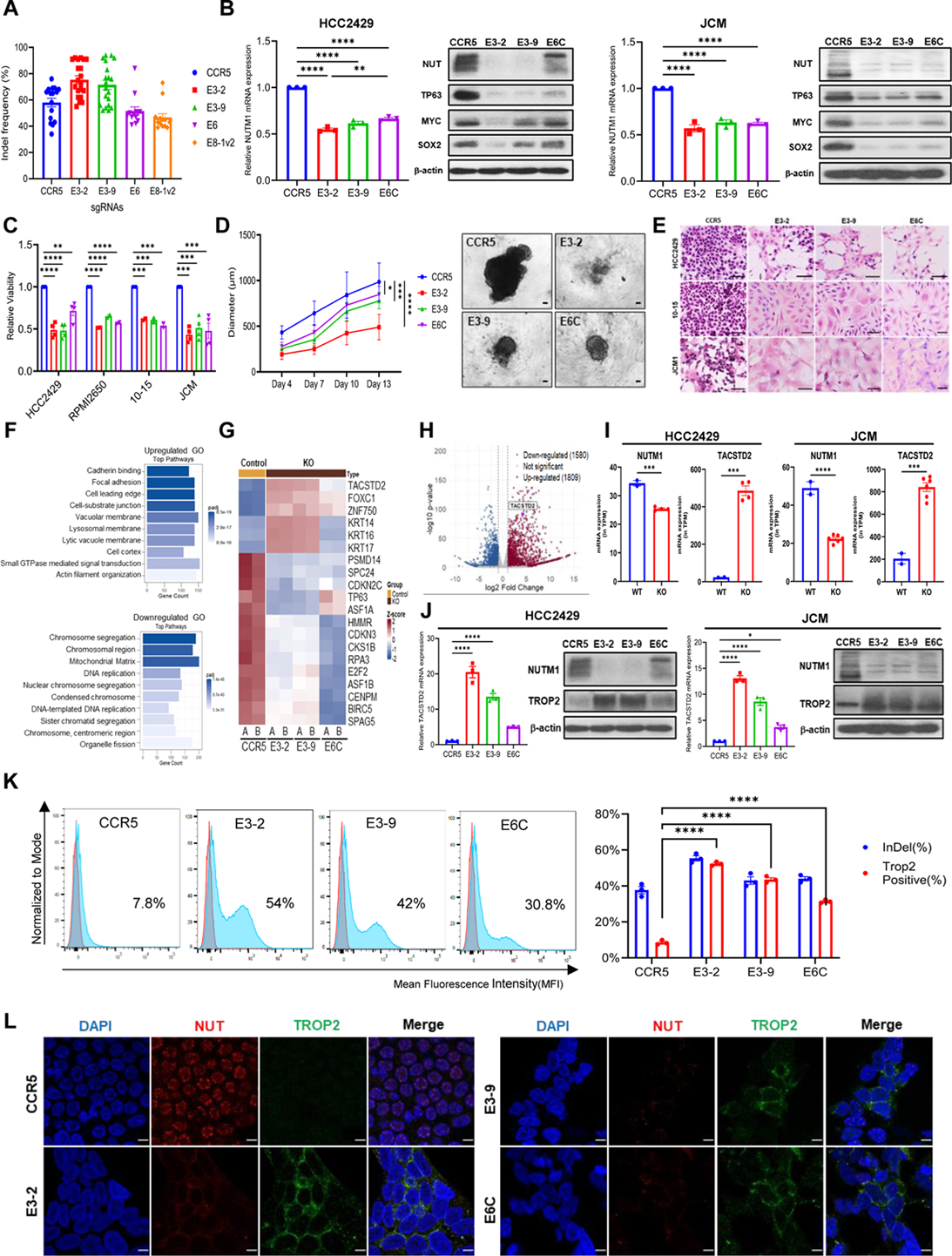
CRISPR-Mediated NUTM1 Suppression Inhibits Tumor Growth and Promotes Differentiation, Concurrent with TROP2 Upregulation in NUT Carcinoma. (A) Quantification of editing efficiency of each sgRNA in the HCC2429 cell line. (B) NUTM1 mRNA expression and western blot analysis of NUT, TP63, MYC, and SOX2 levels in HCC2429 and JCM cell lines. The data are shown as mean ± SEM. P values were determined by 1-way ANOVA with Tukey’s multiple comparisons test. (C) Cell proliferation assay performed on four NC cell lines (HCC2429, RPMI2650, 10-15, JCM1). The absorbance was measured over 7 days. (D) 3D spheroid culture comparing the growth of HCC2429 treated with sgRNAs targeting NUTM1 or CCR5. The spheroid size was measured over 13 days. The data are shown as mean ± SEM. P values were determined by 2-way ANOVA with Tukey’s multiple comparisons test. (E) H&E images of NUTM1-edited NC cells, exhibiting morphological changes. Scale bars represent 50 µm. (F) GO analysis of commonly upregulated and downregulated pathways in the NUTM1-edited NC cell line. (G) Heatmap displaying differential expression of key genes involved in differentiation and proliferation across NUTM1-edited and control HCC2429 cells. (H) Volcano plot of differentially expressed genes between CCR5 and NUTM1-edited HCC2429 cell. TACSTD2 gene was denoted as significant. (I) Barplots showing mRNA expression levels of NUTM1 and TACSTD2 in wild-type and NUTM1-edited JCM and HCC2429 cells. Even a slight reduction in NUTM1 expression resulted in a significant increase in TACSTD2 expression. Data are presented as the mean ± SEM. Statistical significance between the 2 groups was assessed using an unpaired Student’s 2-tailed t test. (J) Relative TACSTD2 mRNA expression and TROP2 expression in HCC2429 and JCM1 cell lines. The data are shown as mean ± SEM. P values were determined by 1-way ANOVA with Tukey’s multiple comparisons test. (K) Flow cytometry analysis shows the proportion of NUTM1-edited cells expressing Trop2 compared to control cells. The TROP2-positive cell fractions closely correlated with the observed indel rates in each edited HCC2429 cell. Data are presented as mean ± SEM, by unpaired 2-tailed Student’s t test. (L) Confocal microscopy images illustrating the reduction of NUT speckled nuclear patterns and the increased TROP2 membrane localization in NUTM1-edited HCC2429 cells. Scale bars represent 5 µm.; *p < 0.05, **p < 0.01, ***p < 0.001, ****p < 0.0001.

### Transcriptome analysis identifies TROP2 upregulation following *NUTM1* suppression

NUT suppression reduced the proliferative capacity and promoted the differentiation of NC tumors. To investigate the underlying molecular changes, we performed transcriptome profiling of *NUTM1*-edited NC cells, which revealed over 6,000 differentially expressed genes, including upregulation of differentiation-related pathways and downregulation of cell cycle programs (Fig. 1F, Supplementary Fig. 4A). Specifically, differentiation markers, including *KRT14*, *KRT16*, *KRT17*, and *FOXC1* were significantly upregulated, whereas proliferation-associated genes, such as *CDKN2C* and *E2F2*, were markedly downregulated, particularly in HCC2429 cells (Fig. 1G). Notably, *TACSTD2* was among the most significantly upregulated genes (Fig. 1H, Supplementary Fig. 4B), prompting further validation given its role as the gene encoding TROP2, a clinically relevant target in antibody-drug conjugate therapies.^24–26^ To investigate the relationship between *NUTM1* and *TACSTD2* expression, we examined their RNA levels in *NUTM1*-edited cells. Despite a modest decrease in *NUTM1* mRNA levels, TROP2 transcript levels increased up to 20-fold in HCC2429 and 13-fold in JCM cells, with a corresponding rise in protein expression confirmed by immunoblotting and flow cytometry TROP2-positive fractions reached 54% in cells edited with sgRNA E3-2, correlating with editing efficiency. (Fig. 1I-K). Immunofluorescence further demonstrated loss of nuclear NUT speckles and prominent TROP2 localization on the membrane (Fig. 1L, Supplementary Fig. 4C). Collectively, these findings suggest that TROP2 upregulation is a direct consequence of *NUTM1* editing and may serve as a potential therapeutic target for differentiated NC tumors.

### Synergistic anti-tumor effects of TROP2-IFN-**β** mutein immunocytokine and *NUTM1* targeting in NC cells

CRISPR/Cas9-mediated *NUTM1* editing in NC cells led to reduced proliferation and promoted differentiation, highlighting its therapeutic potential. However, complete suppression of oncogenic activity may not be achievable with gene editing alone due to the possibility of incomplete knockout and residual fusion protein function. As a result, *NUTM1* editing alone was insufficient to fully inhibit NC cell growth. To address this limitation and enhance therapeutic efficacy, we performed high-throughput drug screening to identify candidate agents that could act synergistically with *NUTM1* suppression. Among the compounds screened, ABBV-744, a selective BET inhibitor, and interferon-beta (IFN-β) mutein, a modified form of interferon-beta with enhanced stability or activity, were identified for their specific cytotoxicity in NC cells and lack of activity in the 293FT cell line, which does not express the fusion gene target. (Fig. 2A, Supplementary Fig. 5). Mechanistically, IFN-β is a critical cytokine involved in initiating T cell-mediated immune responses and has demonstrated anti-tumor potential through both direct tumoricidal activity and enhancement of anti-tumor immunity within the tumor microenvironment. However, the clinical application of IFN-β has been constrained by its short serum half-life and dose-limiting systemic toxicity^27,28^. To address these limitations, antibody-cytokine fusion proteins, or immunocytokines, have emerged as a promising therapeutic strategy by enabling targeted delivery of cytokines to tumors while extending serum half-life^29–32^. In line with this approach, we developed a TROP2-IFN-β mutein immunocytokine. This construct genetically fuses a biologically stabilized IFN-β variant to sacituzumab, an antibody that selectively targets TROP2, which is highly expressed in differentiated regions of NC tumors. This construct enables the precise delivery of IFN-β activity to TROP2-positive tumor cells, offering a targeted immunotherapeutic strategy that complements the effects of *NUTM1* editing. Figure 2B presents a schematic representation of this approach and highlights its potential to enhance therapeutic specificity while minimizing off-target effects. First, we evaluated whether fusing IFN-β to the sacituzumab altered its efficacy in unedited HCC2429 cells, which have a very low TROP2 expression level (Fig. 2B, component C). As expected, both the control monoclonal antibodies and sacituzumab showed minimal to no cytotoxicity. However, the sacituzumab-IFN-β mutein inhibited cell growth by approximately 50%, which indicates that IFN-β activity is not affected by fusion to the antibody. The cytotoxic effect of sacituzumab-IFN-β mutein saturated at lower concentrations was comparable with that of IFN-β mutein alone, which suggests that the conjugated antibody enhanced targeting efficiency even in unedited cells with extremely low TROP2 expression (Fig. 2C). Next, we examined the direct cytotoxicity of the sacituzumab-IFN-β mutein in *NUTM1*-edited cells with elevated TROP2 expression (Fig. 2B, component D). In contrast to the parental and *CCR5*-edited cells, *NUTM1*-edited cells exhibited significantly enhanced cytotoxicity. Specifically, cells edited with sgRNAs E3-2 and E3-9 exhibited IC_50_ values of 13.93 and 16.86 pM, respectively (Fig. 2D). This enhanced efficacy directly correlated with the increased TROP2 expression observed in *NUTM1*-edited cells, thus revealing that higher surface TROP2 levels improved targeting and drug delivery. These findings highlight the potential of combining *NUTM1* editing with TROP2-targeted therapies for achieving more effective treatment outcomes. To further explore the therapeutic potential, we investigated antibody-dependent cellular cytotoxicity (ADCC) and hypothesized that TROP2 upregulation in *NUTM1*-edited cells enhances immune-mediated responses (Fig. 2B, component E). HCC2429 cells were treated with antibodies, either alone or fused with IFN-β mutein, and co-cultured with hCD16-overexpressing NK92MI cells (NK92MI-hCD16) for 4 hours. The effects of ADCC were specifically examined in E3-2 edited cells, which exhibited the highest TROP2 expression levels. As expected, ADCC effects were significantly enhanced, with E3-2 edited cells showing approximately 50% cytotoxicity, indicating a 20% increase compared to the *CCR5*-edited control (Fig. 2E). Together, these results demonstrate that *NUTM1* editing sensitizes NC cells to TROP2-targeted immunocytokine therapy, offering a synergistic strategy that combines tumor reprogramming with enhanced immune-mediated cytotoxicity.

**Figure 2.**
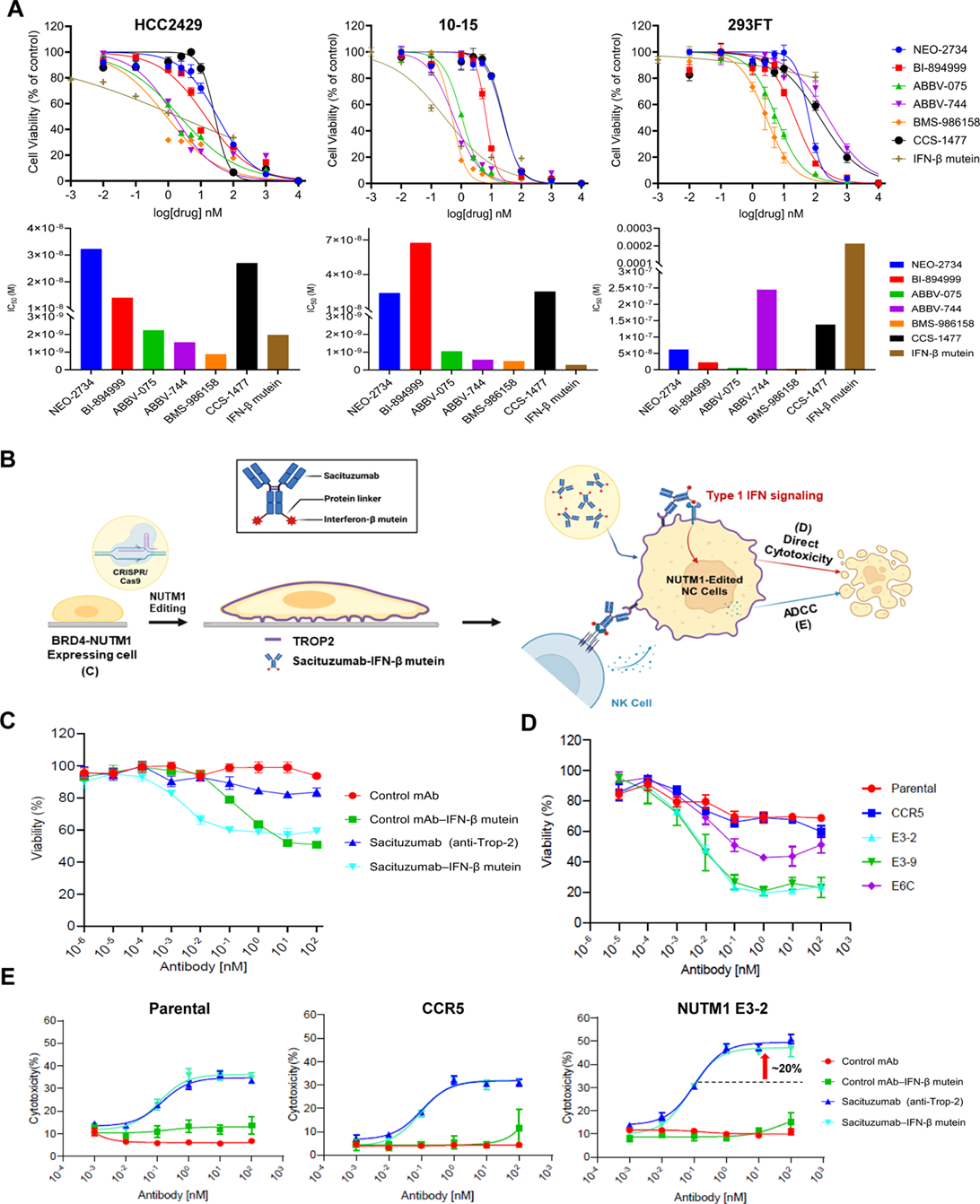
Synergistic anti-tumor effects of TROP2–IFN-β mutein immunocytokine and *NUTM1* targeting in NC cells. (A) Drug screening results across NC cell lines and control 293FT cell line, demonstrating the selective cytotoxicity of IFN-β mutein in NC cells compared with BET inhibitors. (B) Schematic of the combination therapy strategy. *NUTM1* is edited via CRISPR/Cas9, promoting TROP2 expression, followed by treatment with TROP2–IFN-β mutein immunocytokine. This approach leverages both direct cytotoxicity from IFN-β and indirect NK cell-mediated effects. (C) Cytotoxicity assays performed on HCC2429 cells. The cytotoxic effect of sacituzumab–IFN-β mutein saturated at lower concentrations compared with IFN-β mutein alone. (D) Cell viability assay of TROP2–IFN-β mutein ADC treatment following *NUTM1* editing exhibited greater growth inhibition in NC cells compared with controls. (E) NK cell-mediated ADCC sensitivity was increased in NUTM1-edited HCC2429 cells.

### Tumor Differentiation in NC Predicts Clinical Outcomes

Given that NUT suppression led to tumor differentiation and growth inhibition in experimental models, we explored whether this biological effect translates to patient tumors. To assess the clinical implication of tumor differentiation, we analyzed formalin-fixed and paraffin-embedded (FFPE) tumor specimens of treatment-naïve patients. Clinical and pathological features are summarized in Table 1.^33^ Among 27 initially screened cases with positive NUT immunohistochemical (IHC) staining, 24 were confirmed as *NUTM1*-rearranged tumors, characterized by nuclear NUT protein expression in more than 50% of tumor cells (Fig. 3A, Supplementary Fig. 6A).^34^ The remaining three cases with atypical NUT IHC patterns-such as weak cytoplasmic staining or absent nuclear speckles-were excluded following reclassification as SMARCA4-deficient^35,36^ or germ cell tumors (Supplementary Fig. 6B). BRD4 was the predominant fusion partner, detected in 63% of cases, followed by BRD3, NSD3, and YAP1. Fusion partners could not be identified in five cases due to limited RNA quality (Fig. 3B). Histopathologic analysis, based on squamous differentiation, grouped tumors into undifferentiated tumors (UDTs, 50%), poorly differentiated tumors (PDTs, 21%), and differentiated tumors (DTs, 29%) (Fig. 3C). Kaplan-Meier analysis showed significantly better progression free survival (PFS) (p = 0.037) and OS (p = 0.0096) in patients with DTs compared to those with UDTs or PDTs (Supplementary Fig. 7A). These differences were even more pronounced in thoracic NCs harboring bromodomain-containing fusion partners, with PFS and OS p-values of 0.019 and 0.0026, respectively (Fig. 3D).

**Figure 3.**
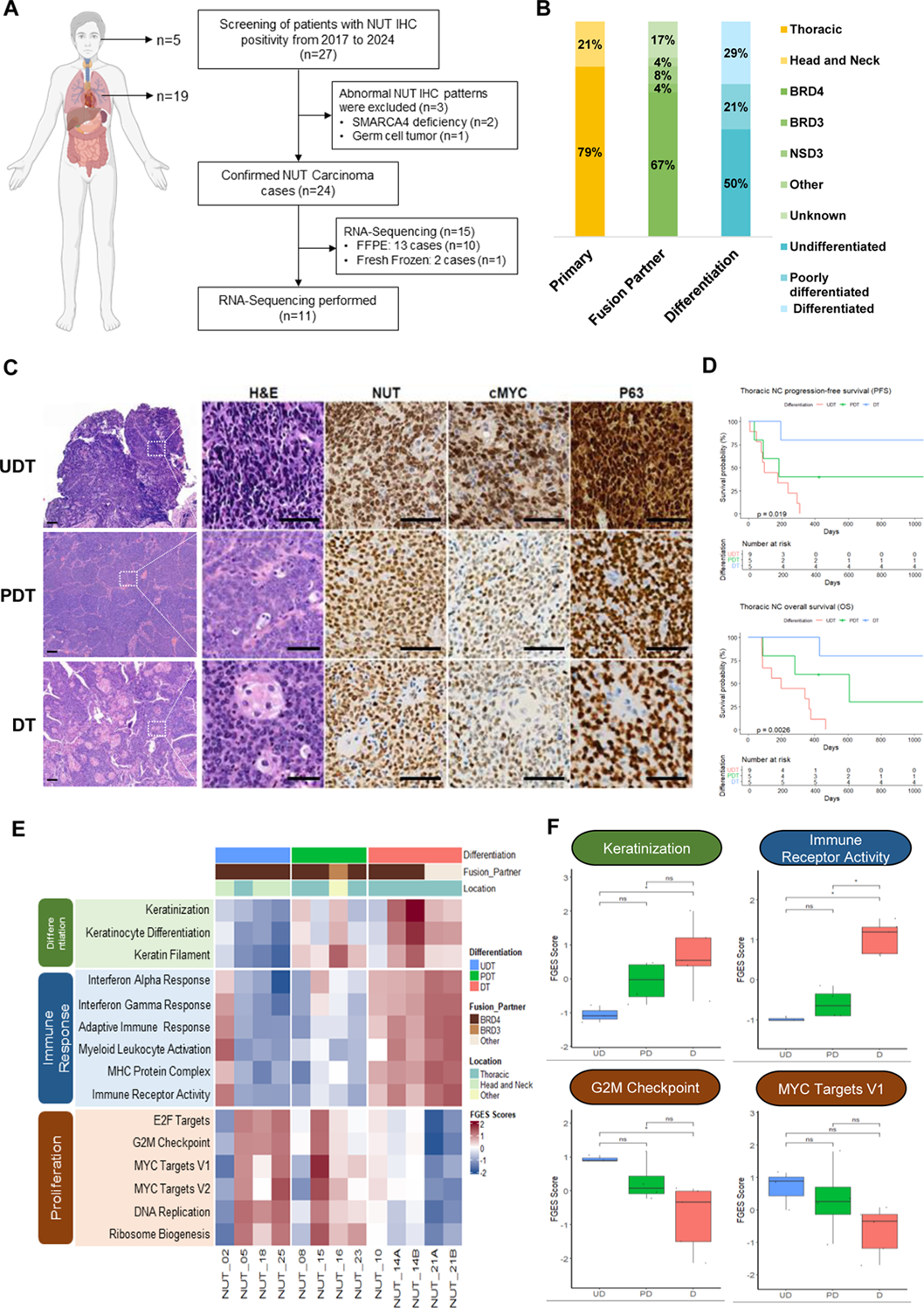
Integrated Histologic, Molecular, and Clinical Profiling Defines Differentiation-Dependent Subgroups in NUT Carcinoma. (A) Schematic representation of the anatomical distribution of patients with NC and the number of patient samples analyzed by RNA sequencing. (B) Distribution of primary tumor location, fusion partners, and differentiation status in patients with NC. (C) Histopathological classification of NC tumors into UDT, PDT, and DT groups with representative H&E and IHC images of NUT, c-MYC, and p63. DTs exhibit squamous pearls. (D) Kaplan–Meier survival curve of PFS and OS stratified by tumor differentiation status of patients with Thoracic NC. (E) Heatmap of functional gene enrichment scores (FGES) for pathways categorized into proliferation, differentiation, and immune response. (F) Boxplots showing FGES for representative pathways. Statistical significance was determined using the Kruskal–Wallis test followed by the Wilcoxon rank-sum test (*p < 0.05, **p < 0.01, and ***p < 0.001).

**Table 1.** Clinical characteristics of NUT carcinoma cases.

| Case No. | Sex | Age (yr) | Primary Location | Metastasis | Treatment | Partner | Differentiation | Trop2 | Survival | OS (mo) |
| --- | --- | --- | --- | --- | --- | --- | --- | --- | --- | --- |
| 1* | M | 45 | Lung | Yes | CRT | BRD4(E11) | DT | Pos. | Yes | 84.5 |
| 2* | M | 43 | Nasal cavity | No | S + CRT | BRD4(E11) | UDT | Neg. | No | 14.0 |
| 3* | M | 32 | Lung | Yes | CT | NSD3 (E7) | UDT | Pos. | No | 2.8 |
| 4* | M | 34 | Lung | Yes | CRT | N/A | UDT | N/A | No | 12.4 |
| 5* | F | 74 | Trachea | Yes | CRT | BRD4(E11) | UDT | Neg. | No | 11.2 |
| 6* | M | 37 | Bronchus | Yes | S + CT | NSD3 (E7) | DT | Pos. | Yes | 71.4 |
| 7* | M | 49 | Pleura | Yes | S + CRT | BRD4(E14) | UDT | Pos. | No | 2.7 |
| 8* | F | 41 | Pleura | Yes | CT | BRD4(E11) | PDT | Pos. | No | 2.9 |
| 10* | F | 48 | Lung | No | S + CT | BRD4(E11) | DT | Pos. | Yes | 57.8 |
| 11* | M | 43 | Lung | No | S + CT | BRD4(E11) | DT | Pos. | Yes | 53.1 |
| 12* | F | 18 | Bronchus | Yes | S + CRT | BRD4(E11) | PDT | Pos. | No | 20 |
| 13* | M | 44 | Bronchus | Yes | CRT | BRD4(E11) | UDT | Neg. | No | 2.8 |
| 14* | F | 50 | Lung | No | S + CRT | BRD4(E11) | DT | Pos. | No | 14.1 |
| 15* | F | 72 | Pleura | Yes | S | BRD4(E10) | PDT | Neg. | No | 9.3 |
| 16* | M | 49 | Lung | No | CRT | BRD3(E9) | PDT | Pos. | Yes | 43.3 |
| 17* | M | 44 | Lung | Yes | CRT | BRD4(E11) | UDT | N/A | No | 15.4 |
| 18 | M | 36 | Lung | Yes | S + CRT | BRD4(E10) | UDT | Neg. | No | 4.7 |
| 19 | M | 34 | Lung | Yes | CRT | N/A | UDT | N/A | No | 6.5 |
| 20 | M | 29 | Submandibular Gland | Yes | S + CRT | N/A | UDT | Neg. | Yes | 23.9 |
| 21 | F | 68 | Lung | No | S | YAP1(E3,4) | DT | Pos. | Yes | 20.5 |
| 23 | M | 25 | Lung | No | S | BRD4(E13) | PDT | Neg. | Yes | 14 |
| 24 | F | 62 | Lung | Yes | CRT | N/A | UDT | Neg. | No | 12 |
| 25 | M | 68 | Nasal cavity | No | S + CRT | BRD4(E14) | UDT | Neg. | Yes | 8 |
| 27* | M | 70 | Supraglottic | No | S | BRD4(E14) | UDT | Neg. | Unknown | Unknown |

M, Male; F, Female; S, Surgery; CT, Chemotherapy; CRT, Chemotherapy+Radiotherapy; UDT, Undifferentiated Tumor; PDT, Poorly Differentiated Tumor; DT, Differentiated Tumor; yr, year; mo, month
\*, Previously reported cases at Samsung Medical Center<sup>33</sup>

### Differentiation-Associated Transcriptomic Signatures and Immune Activation

To investigate the molecular signatures associated with differentiated tumor cell populations within NC tumors, we further analyzed their transcriptomic profiles. Principal component analysis demonstrated clear separation of DTs from UDTs and PDTs, indicating distinct global expression profiles (Supplementary Fig. 7B). Given the known role of MYC in NC aggressiveness, we compared MYC pathway activity across differentiation states and observed a marked reduction in DTs relative to PDTs and UDTs (Supplementary Fig. 7C). We also identified biological distinctions among the differentiation subgroups through gene set variation analysis (GSVA). PDTs and UDTs were enriched for proliferative signatures, such as G2M checkpoint, E2F targets, and MYC targets, consistent with their more aggressive clinical behavior. In contrast, DTs exhibited downregulation of cell cycle-related pathways, including DNA replication and ribosome biogenesis, reflecting reduced proliferative capacity (Fig. 3E, Supplementary Fig. 7D). Notably, immune-related pathways-particularly immune receptor signaling and interferon responses-were significantly upregulated in DTs. These findings suggest that an active immune microenvironment may contribute to the improved prognosis observed in differentiated tumors. In parallel, DTs showed higher expression of keratinization and keratinocyte differentiation genes, further supporting the association between transcriptional and histological differentiation (Fig. 3F). These observations, consistent with our *in vitro* finding, demonstrate that tumor differentiation status is closely associated with both proliferative activity and immune signaling in NC. DTs exhibit reduced oncogenic programs and enhanced immune activity, aligning with their favorable clinical outcomes.

### Spatial Analysis of Differentiation-Associated Molecular and Immune Features

Spatial transcriptomic profiling was performed on a treatment-naïve NUT carcinoma sample (patient NUT14) to characterize the spatial basis of transcriptional heterogeneity observed in RNA-seq. The tumor exhibited histologic heterogeneity, with distinct PDT and DT regions. This case provided a unique opportunity to directly compare tumor microenvironments within the same lesion across differentiation states. Specimen characteristics, including gross features, tumor size, and TNM stage, are summarized in Supplementary Figure 8.

Spatial annotation was performed manually based on histologic features (see Supplementary Fig. 8 and supplementary materials). Based on these annotations, tumor and stromal regions were annotated into PDT, DT, PDT-DT mixed, and stromal subtypes (Fig. 4A). Uniform manifold approximation and projection (UMAP) clustering revealed spatial segregation between PDT and DT regions, with DTs located adjacent to lymphocyte-enriched stroma, suggesting enhanced immune interactions (Fig. 4B). Cell-of-origin inference using SingleR^37^ confirmed this distinction, with PDTs aligning with LUSC and LUSC mitotic profiles, and DTs annotated as mixed NSCLC subtypes (Fig. 4C-D, Supplementary Fig. 8H-J). These findings not only support recent proposals to reclassify NUT carcinoma as a subtype of SCC^38^, but also suggest that transcriptional plasticity may underlie the emergence of intratumoral heterogeneity through differentiation.

**Figure 4.**
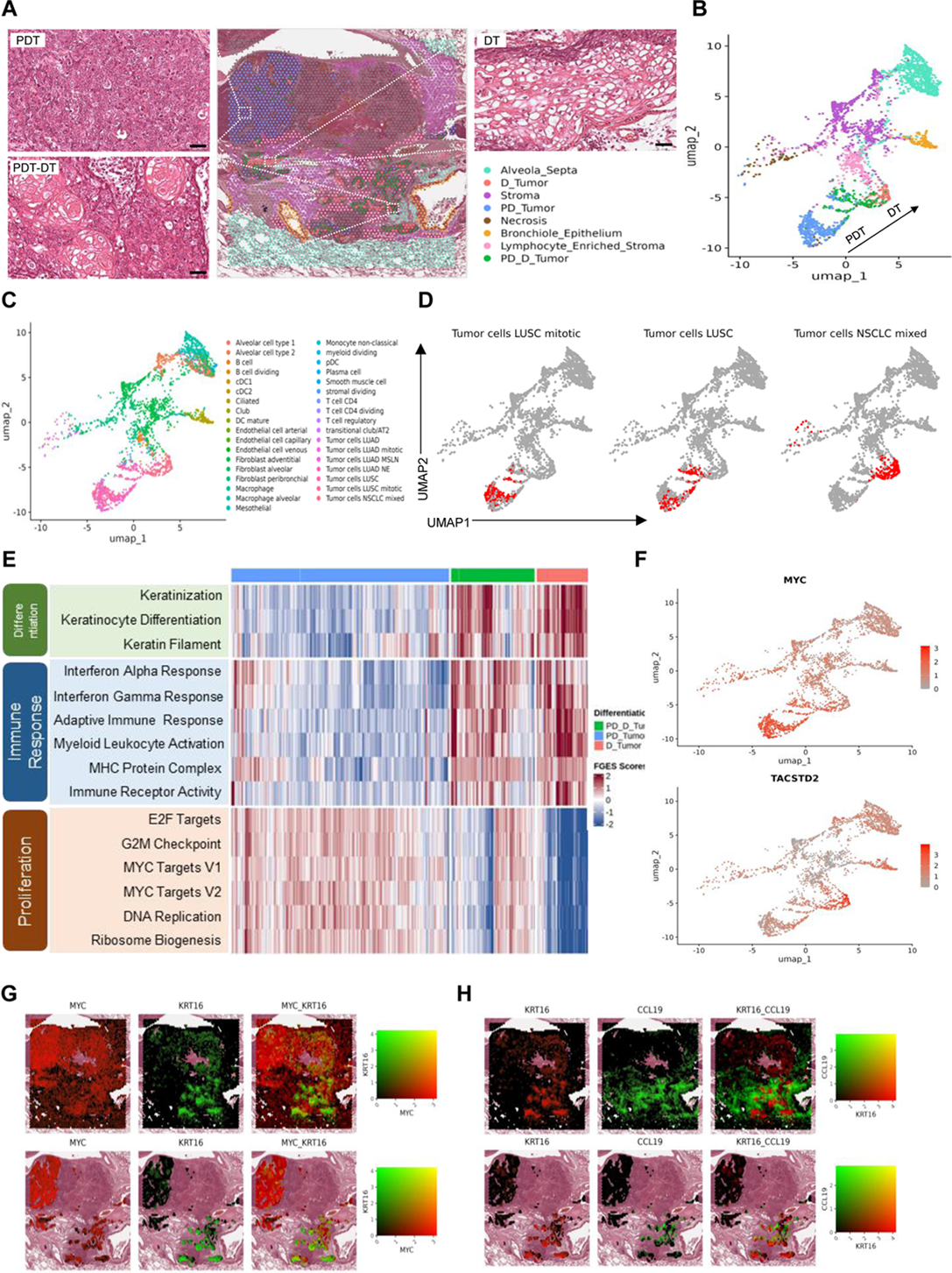
Tumor Differentiation in NUT Carcinoma Associates with Distinct Spatial Immune and Proliferative Profiles. (A) Manually annotated spatial transcriptomics of a treatment-naïve NUT carcinoma patient (NUT14), highlighting tumor subregions and the surrounding microenvironment. Scale bar, 50 μm. (B) UMAP clustering reveal that differentiated tumor regions are spatially closer to lymphocyte-enriched stroma, suggesting enhanced immune interaction. (C) singleR based cell of origin analysis using the LuCA single-cell lung cancer dataset validates manual annotations. (D) Cell of origin analysis further classifies PDT regions as squamous carcinoma subtypes, while DT regions map to NSCLC-mixed subtypes. (E) Heatmap of FGES from gene set variation analysis results of the PDT, PDT–DT, and DT pathways. Pathways are categorized into proliferation, differentiation, and immune responses. (F) Differential spatial expression of MYC and TACSTD2 distinguishes proliferative from differentiated tumor areas. (G) Spatial feature plots demonstrate mutually exclusive expression of MYC and KRT16, indicating that differentiation is inversely associated with proliferation. (H) Spatial co-localization of KRT16 and CCL19 suggests that differentiated regions may exhibit enhanced immune activation.

Spatial transcriptomic analysis revealed proliferative signatures in PDT regions, including G2M checkpoint, E2F targets, and MYC pathways, while DT regions showed enrichment of immune-related pathways such as adaptive immune and interferon responses (Fig. 4E, Supplementary Fig. 8K-N). MYC expression was localized to PDTs, whereas *TACSTD2* was enriched in DTs (Fig. 4F). Co-localization analyses further illustrated mutually exclusive expression of proliferative (MYC) and differentiation markers (KRT4, KRT16, SCEL), alongside DT-specific expression of immune genes including IGHA1, IGHG1, and CCL19 (Fig. 4G, Supplementary Fig. 8O). Notably, co-expression of KRT16 and CCL19 within DTs suggests a possible association between tumor differentiation and immune activation (Fig. 4H, Supplementary Fig. 8P-Q).

Collectively, these spatial transcriptomic data reveal molecular and immune heterogeneity in NC and demonstrate that DT regions are associated with immune-enriched microenvironments and reduced proliferative activity, whereas PDT regions display an immune-cold, proliferative profile. These findings validate RNA-seq results and highlight tumor differentiation as an important factor influencing tumor aggressiveness and immunogenicity in NC, supporting the rationale for differentiation-based therapeutic strategies.

### TROP2 expression correlates with differentiation and favorable clinical outcomes in NC

We next assessed TROP2 expression in NC tumors to evaluate its association with tumor differentiation status and clinical outcomes. TROP2 positivity was most frequent in DTs (100%), followed by PDTs (60%) and UDTs (22.2%), showing a strong positive correlation with *TACSTD2* mRNA levels (Spearman’s ρ = 0.664, p = 0.001; Fig. 5A-D). TROP2 expression was observed in regions containing differentiated tumor cells, suggesting a correlation between TROP2 expression and tumor differentiation. This also indicates that a subset of NC tumors exhibits a heterogeneous tumor cell population. To evaluate the association between TROP2 expression and clinical outcomes, we performed Kaplan-Meier survival analyses. Although not statistically significant, patients with TROP2-positive tumor cells exhibited a trend toward longer OS (median OS: 20.5 vs. 11.2 months; p = 0.19) and patients with metastatic disease had significantly worse survival outcomes (p = 0.013; Fig. 5E). We further conducted a Cox proportional hazards analysis to evaluate potential prognostic factors, including differentiation status, presence of metastasis, and fusion gene partners. DTs were associated with a significantly lower hazard ratio (HR) for death compared to UDTs (HR = 0.08; 95% confidence interval [CI], 0.010-0.61; p = 0.02). Conversely, the presence of metastasis was associated with a significantly increased risk of death (HR = 3.83; 95% CI = 1.230-11.90; p = 0.02) (Fig. 5F). Together, these findings highlight that TROP2 expression is closely associated with tumor differentiation and may serve as a favorable prognostic marker in NC, supporting its potential utility in both tumor characterization and risk stratification.

**Figure 5.**
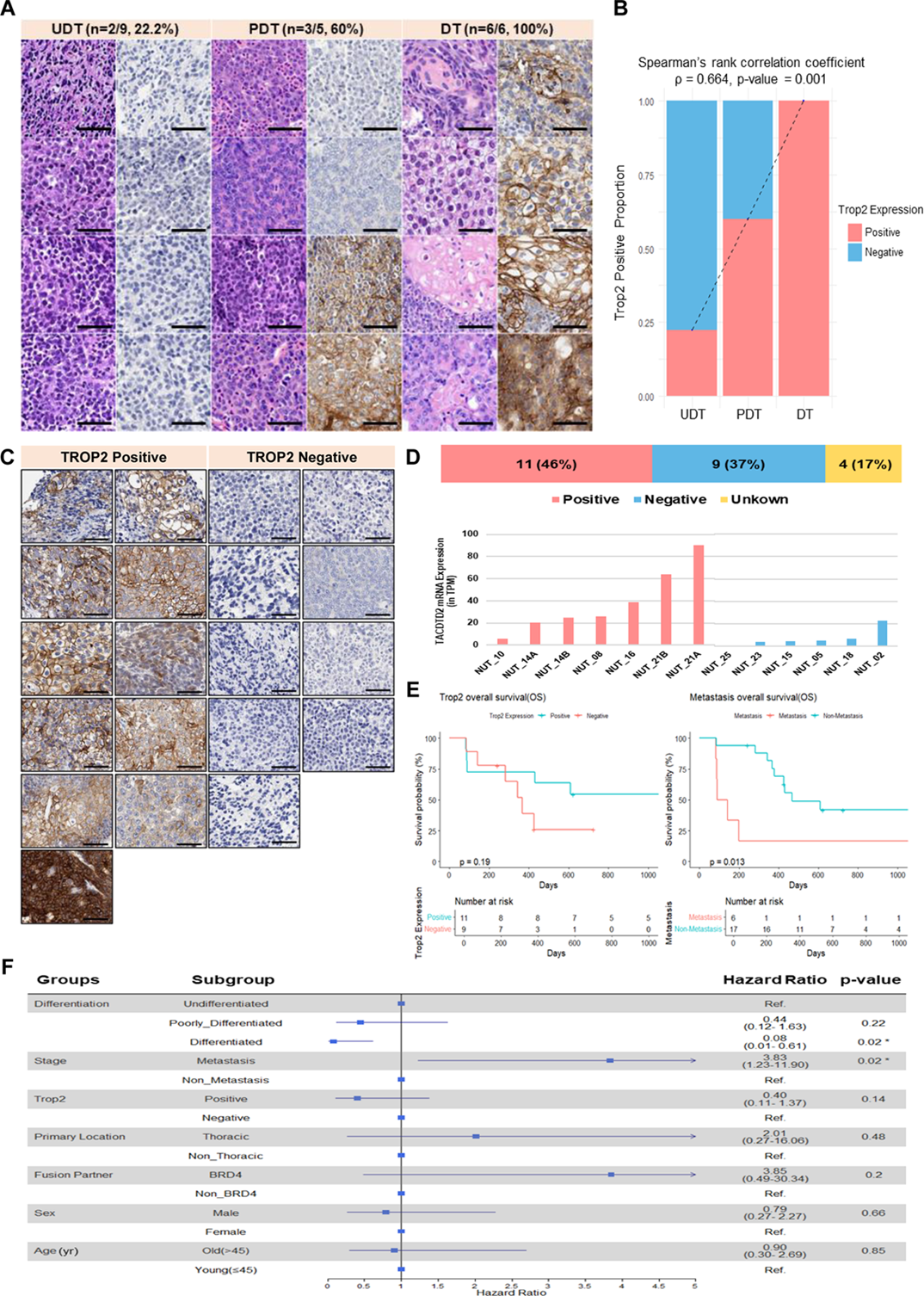
Exploring the link between TROP2 expression and survival in patients with NUT carcinoma. (A) Representative images of H&E and TROP2 IHC staining across different differentiation statuses in samples from patients with NC. Increased TROP2 positivity was observed in the more differentiated tumors. Scale bar is 50µm. (B) Correlation between TROP2-positivity and the differentiation status was determined using Spearman’s correlation coefficient. (C)IHC images showing TROP2 staining in NC patient tumor samples. Scale bars represent 50 µm. (D) Distribution of TROP2 expression in the NC cohort. The degree of TROP2 staining generally correlates with *TACSTD2* mRNA expression. (E) Kaplan–Meier survival analysis based on TROP2 expression. Patients with TROP2-positive tumors exhibit improved median OS than the patients with TROP2-negative tumors. (E) Cox proportional hazards model showing the impact of clinical variables on OS. Differentiated tumors and absence of metastasis were associated with better survival outcomes.

**Figure 6.**
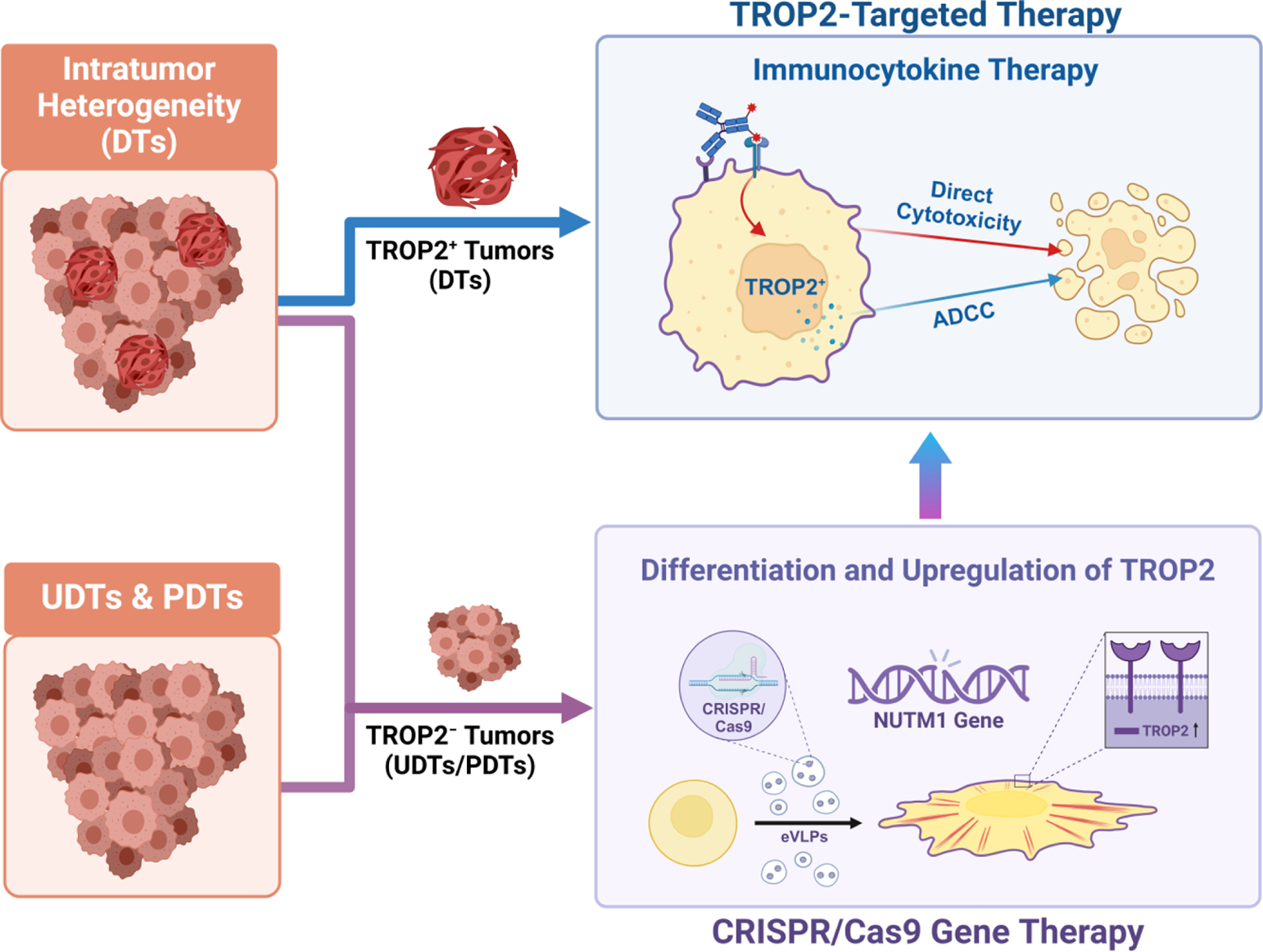
Graphical summary. This graphical summary illustrates a sequential targeting therapeutic strategy for NC, addressing the heterogeneity within NUT carcinoma by tailoring treatments based on TROP2 expression and differentiation status and ensuring that all tumor subtypes can be effectively targeted.

## Discussion

This study introduced a novel sequential targeting strategy for NC that combines CRISPR/Cas9-mediated NUT suppression with TROP2-targeted immunocytokine therapy. Unlike BETis, which non-selectively targets the bromodomains of *NUTM1* fusion partners and exhibits limited efficacy and substantial adverse effects, NUT suppression offers a more precise and tumor-specific approach. By targeting *NUTM1* directly, this strategy suppresses tumor growth and induces differentiation, thereby altering tumor phenotypes and enhancing therapeutic responsiveness. Our transcriptomic analysis revealed distinct molecular profiles associated with tumor differentiation in NC. DTs upregulate immune-related pathways, including complement and interferon responses and downregulate proliferative pathways, including MYC signaling. In contrast, PDTs and UDTs exhibit reduced immune activity and elevated proliferative signaling, aligning with their more aggressive phenotypes and poorer clinical outcomes. Most long-term survivors (>2 years) exhibited DT regions and TROP2 positivity (Table 1), which further supports the association between differentiation and a favorable prognosis. Therefore, NUT suppression not only induced differentiation but also upregulated TROP2 expression, sensitizing tumors to TROP2-IFN-β mutein immunocytokine therapy and enhancing immune-mediated cytotoxicity. DTs with high TROP2 expression benefit from TROP2-IFN-β mutein immunocytokine therapy owing to synergistic effects between immune activation and the therapeutic mechanism of action. For PDTs and UDTs with low TROP2 expression, CRISPR/Cas9-mediated *NUTM1* editing offers a strategy for inducing differentiation, upregulating TROP2, and improving the responsiveness to targeted therapies. This study proposes that a sequential targeted treatment, grounded in the principles of targeted therapy, accurately captures the molecular characteristics of tumors, thereby enhancing treatment specificity and efficacy, and ultimately enabling effective therapeutic responsiveness.

Despite extensive research on the genetic and molecular features of NC^17,39^, the prognostic significance of tumor differentiation remains relatively understudied. A previous study suggested a potential association between specific histological characteristics and clinical outcomes; however, the limited sample size reduced the statistical power, preventing definitive conclusions^3^. In this study, tumors exhibiting focal keratinization or squamous pearls were classified as differentiated, enabling us to demonstrate that the presence of differentiated cells may serve as a meaningful prognostic indicator in NC. The discrepancy across studies may be attributed to differences in the classification criteria, and our more nuanced categorization may allow for a more precise distinction of the differentiation status and its role as a meaningful prognostic marker for NC. Transcriptome analysis revealed a strong association between differentiation status, immune activity, and prognosis in NC; however, the causal relationship between differentiation and immune activation remains unclear. In gastric cancer (GC), poorly differentiated cells exhibit reduced immune activity, whereas differentiated cells are associated with an immune-rich environment and enhanced immune cell infiltration. The process of trans-differentiation in GC, where differentiated cells transform into neuroendocrine carcinoma cells, involves reduced immune infiltration and downregulation of immunostimulatory genes, indicating a dynamic interplay between differentiation and immune response^40,41^. Applying these insights to NC, our findings revealed that DTs upregulated immune pathways, such as complement and interferon responses, fostering an immune-active microenvironment conducive to therapeutic engagement. Spatial transcriptomic data further supported this observation, showing the co-localization of *TACSTD2* and immune-related genes in DT regions, whereas MYC, a proliferation marker, was predominantly expressed in PDTs (Supplementary Fig. 8Q). These findings suggest that inducing tumor differentiation in NC may promote an immune-active tumor microenvironment, potentially establishing a positive feedback loop that contributes to less aggressive tumor behavior and improved clinical outcomes. This hypothesis is supported by the observation that TROP2-positive, differentiated tumors are predominantly found in long-term survivors.

Nevertheless, not all cases align consistently with this proposed model. There were molecular inconsistencies observed in tumors classified as UDT. Patient NUT02, categorized as having UDT with negative TROP2 IHC staining, exhibited relatively high *TACSTD2* mRNA expression and low proliferation marker levels, which suggests heterogeneity within the UDT group. This discrepancy may be explained by the presence of normal tissue regions (approximately 20%) within the tumor sample, wherein signals from normal cells disproportionately influenced the transcriptomic profile (Fig. 3E, 5D). Furthermore, the rarity of NC inevitably constrained cohort size, limiting the statistical power needed to robustly assess correlations between TROP2 expression and tumor differentiation. To more precisely investigate differentiation-associated intratumoral heterogeneity in actual patient tissues, we employed spatial transcriptomic analysis to visualize the tumor microenvironment (Fig. 4). This approach enabled us to explore previously underappreciated aspects of NC, such as differentiation-associated heterogeneity and its relationship with the tumor immune landscape. Although the underlying mechanisms remain unclear, our data reveal the presence of poorly differentiated tumor cells within NC that appear to initiate immune responses and promote differentiation. This trans-differentiation process is associated with reduced tumor aggressiveness and increased expression of TROP2, suggesting a potential link between immune activation, differentiation, and therapeutic responsiveness.

Although our study has certain limitations inherent to a proof-of-concept investigation, it provides important insights into the therapeutic potential of a sequential targeted strategy for NC, an ultra-rare malignancy with limited treatment options. While the *in vitro* data strongly supports the proposed approach, *in vivo* validation remains challenging due to the lack of physiologically relevant NC models. Most available models are derived from highly proliferative, undifferentiated tumor components, introducing a selection bias that limits the study of differentiation dynamics. Genetically engineered mouse models offer unique advantages in modeling tumor-immune interactions, but remain underdeveloped for *NUTM1* fusions. In particular, achieving tissue- and stage-specific expression to mimic the disease context is technically demanding. Together, these limitations highlight the need for advanced models that better recapitulate the interplay between differentiation, TROP2 expression, and immune activation in NC.

Nevertheless, our study is significant in that it proposes an advanced, mechanism-based therapeutic framework for a molecularly defined but clinically underserved cancer. By addressing tumor heterogeneity and identifying actionable additional targets such as TROP2, these findings support the clinical relevance of pursuing translational studies and developing precision-based therapeutic strategies in NC. In conclusion, this study introduces a novel sequential targeting strategy for NC, combining CRISPR/Cas9-mediated NUT suppression with TROP2-targeted immunocytokine therapy. This approach suppresses tumor growth, induces differentiation, and upregulates TROP2 expression, thereby fostering an immune-active microenvironment and enhancing therapeutic specificity. Differentiation has been highlighted as a potential prognostic marker and therapeutic target in NC and is closely linked to immune activation and improved clinical outcomes. Future studies should validate these findings in larger cohorts and *in vivo* models, emphasizing the mechanisms regulating TROP2 expression and clarify the causal dynamics between differentiation and immune activation. Such efforts could refine therapeutic strategies and translate this approach into effective clinical applications and ultimately improve the treatment outcomes of patients with NC.

## Methods

### Cell culture

The NC cell lines SNU-2972-1, SNU-3178S, Ty-82, and HCC2429 were generously provided by Prof. Kim at the Seoul National University Hospital, Seoul, South Korea. The JCM and 10-15 NC cell lines were kindly provided by Prof. Christopher A. French from Boston, USA. RPMI2650 cells were obtained from the Korean Cell Line Bank (#10030). SNU-2972-1, SNU-3178S, Ty-82, RPMI2650, and HCC2429 cells were cultured in RPMI-1640 medium (Cytiva, #SH30027.01) supplemented with 10% fetal bovine serum (Gibco, #16000-044) and 1% penicillin–streptomycin–amphotericin B (Thermo Fisher Scientific, #15240-062). JCM and 10-15 cells were cultured in Dulbecco’s modified eagle medium (Cytiva, #SH30243.01) under the same conditions. All NC cells were cultured at 37 °C with 5% carbon dioxide and were confirmed to be negative for mycoplasma by testing with a MycoFluor™ Mycoplasma Detection Kit (Thermo Fisher Scientific, #M7006). The cells were authenticated via short-tandem repeat profiling at Macrogen.

### 3D spheroid culture

Aliquots of 50-, 100-, and 250-edited HCC2429 cells were plated in 96-well round cell floater plates (SPL Life Sciences, #34896) to assess growth, and both the long- and short-axis diameters were measured.

### sgRNA design

sgRNAs targeting the *NUTM1* gene for CRISPR/Cas9 were designed using the CRISPick online software. Human GRCh37 or GRCh38 was used as the reference genome, with the CRISPRko method and the SpCas9 (NGG) enzyme set to default. sgRNAs were selected based on their high specificity rankings and low potential off-target scores. Additional sgRNAs, including Sg E3-6, 3-7, 3-8, and 3-9, were designed manually without using design software. The first nucleotide of the spacer sequence was ensured to be “G” to facilitate cloning. Spacer sequences of four consecutive nucleotides were excluded to prevent potential complications. Furthermore, sequences with minimal similarity to other genomic sequences were selected and confirmed using Cas-Offinder.

### Cas9-eVLP production

Cas9-eVLPs were produced via transient transfection of Lenti-X 293T cells (Clontech, #632180). Cells were seeded at a density of 1 × 10^7^ cells per 150-mm dish. After 24 h, the cells were co-transfected with VSV-G (800 ng; Addgene, #12259), gag-pol (6750 ng; Addgene, #35614), gag-Cas9 (2250 ng; Addgene, #181752), and sgRNA (8800 ng) using 120 μg of polyethylenimine (Polyscience Inc., #23966-100). Subsequently, 48 h after transfection, the supernatant was collected and centrifuged at 500 *g* for 5 min to remove the cellular debris. The supernatant containing eVLPs was then filtered through a 0.45-μm CA filter (Sartorius, #S6555-FMOSK). To precipitate the eVLPs, 5× PEG-it Virus Precipitation Solution (System Biosciences, #LV825A-1) was added to the supernatant and incubated overnight at 4 °C. On the next day, eVLPs were pelleted via centrifugation at 1,500 *g* for 30 min at 4 °C and concentrated by resuspending in 1× HIV-safe Manager (Lugen SCI, #LGV- 1022B). The eVLPs were stored at -80 °C and thawed on ice immediately before use. To ensure consistency across experiments, all eVLPs used in the cell culture studies were concentrated using this standardized method^42^, allowing for direct comparison of Cas9-eVLP potency at equal volumes. This concentration method yielded approximately 6–6.5 × 10^8^ eVLPs/μL.

### Quantification and titer determination of Cas9-eVLPs

The concentration of MLV p30 protein in the Cas9-eVLPs was measured using a MuLV Core Antigen ELISA kit (Cell Biolabs, #VPK-156), following the manufacturer’s instructions. To estimate the concentration of VLP-associated p30 protein, 20% of the p30 detected in the solution was assumed to be associated with VLPs. The number of Cas9 protein molecules per eVLP was calculated by assuming that each eVLP contained approximately 1800 molecules of p30, based on earlier reports on MLV particles^43^. This method was used to determine the eVLP titers.

### Measurement of mutation frequency

#### eVLP transduction and genomic DNA extraction

Cells were seeded in 24-well plates (30048, SPL) at a density of 1–2.5 × 10^5^ cells per well, depending on the cell line. After 20–24 h, Cas9-eVLPs were added directly to the culture medium in each well. At 72 h after transduction, the cells were harvested, and genomic DNA was extracted using a DNA Mini Kit (Qiagen, #51306) following the manufacturer’s instructions. The extracted DNA was quantified using a NanoDrop spectrophotometer. PCR amplification was conducted using the AccuPower Hotstart PCR premix (Bioneer, #K5051-1) with primers specific to the target sequences.

#### T7E1 cleavage assay

PCR amplicons from the targeted genomic region were purified using the QIAquick PCR Purification Kit (Qiagen, #28106). The purified PCR products were denatured and annealed in NEBuffer 2 (NEB, #B7002S) using a thermocycler. Hybridized PCR products were digested with T7 endonuclease 1 (NEB, # M0302L) for 30 min at 37 °C and subjected to 2% agarose gel electrophoresis. Supplementary Table 3 enlists all PCR primer sequences.

#### ICE analysis

Genome editing efficiency was assessed using the Synthego online tool Interference of CRISPR Edits (ICE) Analysis. The ICE analysis tool identifies major induced mutations at the editing site and quantifies their frequency within the cell population based on Sanger sequencing data from both parent and gene-edited cell lines.

#### Targeted deep sequencing

The extracted gDNA was used to amplify the target site using PCR to construct a sequencing library. The PCR conditions were as follows: 95 °C for 3 min; 30–35 cycles of 95 °C for 30 s, 61 °C for 30 s, and 72 °C for 30 s; followed by 72 °C for 5 min. The PCR products were purified and sequenced using iSeq 100 (Illumina Inc.). The mutation frequency was analyzed using the MOUND software (available at https://github.com/ibs-cge/maund).

### qRT-PCR assays

RNA was extracted and quantified as described previously. cDNA was synthesized from 1 μg of total RNA using SuperScript™ IV Reverse Transcriptase (Thermo Fisher Scientific, #18090200) following the manufacturer’s protocol. qRT-PCR was performed on either the PRISM® 7900HT Fast Real-Time PCR System (Applied Biosystems) or QuantStudio™ 6 Flex Real-Time PCR System (Applied Biosystems) using 2× SYBR Green Master Mix (Applied Biosystems, #4368702) in 384-well plates. Gene expression was normalized to that of the housekeeping gene GUSB. Supplementary Table 3 enlists the primer sequences used in these assays. All experiments were conducted in triplicate for each sample, and statistical analyses were performed using unpaired Student’s two-tailed t-test.

### Cell viability assay

Transfected cells were seeded into 96-well plates at a density of 500–5,000 cells per well in a total volume of 100 μL media, depending on the cell line. The cell viability was evaluated at various time points based on the purpose of the experiment. The culture medium in each well was replaced with a mixture of 10 µL of Ez-Cytox (DoGenBio, #EZ-3000) and 90 µL of fresh culture medium. The plates were incubated at 37 °C in a humidified atmosphere containing 5% CO_2_ for 2–4 h. Absorbance was measured at 450 nm using a SpectraMax® 96-well plate reader (Molecular Devices).

### Immunohistochemistry

Immunohistochemical analysis was performed for all cases using 4-μm-thick serial sections from a representative FFPE block. All protein-antigen analyses were performed using a Ventana BenchMark XT automatic immunostainer, following the manufacturer’s instructions (Ventana Medical Systems).

### Immunoblotting

Cells were lysed in 1× RIPA buffer with additional NaCl to a final concentration of 250 mM (Cell Signaling Technology, #9806S) containing protease inhibitors at 4 °C for 15 min with rotation. The lysates were then centrifuged at 13,000 rpm for 25 min at 4 °C, and the supernatants were collected for protein quantification using either the BSA or Bradford assay. Protein lysates were separated using sodium dodecyl sulfate polyacrylamide gel electrophoresis and transferred onto polyvinylidene fluoride membranes (Millipore, #IPVH00010). The membranes were blocked with 5% skim milk for 30 min, followed by an overnight incubation at 4 °C with gentle rocking in the presence of primary antibodies. Subsequently, the membranes were washed twice with Tris-buffered saline with 0.1% Tween® 20 detergent (TBST) at room temperature for 10 min each, followed by a 1-h incubation with secondary antibodies diluted 1:5,000 in 5% skim milk at room temperature. The membranes were then washed three times for 10 min each in TBST, developed using an electrochemiluminescence reagent (GE Healthcare, #12316992), and visualized by exposing to a film (AGFA, #CP-BU). Supplementary Table 4 enlists the antibodies used.

### Immunofluorescence staining and confocal microscopy

Coverslips with cultured cells were washed with phosphate buffered saline (PBS) and fixed with 4% paraformaldehyde for 10 min, followed by three PBS washes. The coverslips were then incubated in blocking buffer for 1 h. After removing the buffer, primary antibodies were applied and incubated overnight at 4 °C. On the next day, the coverslips were washed three times with PBS for 5 min. Thereafter, fluorescently labeled secondary antibodies were applied and incubated for 1–2 h in the dark. Next, the coverslips were washed three times with PBS for 5 min each in the dark. The coverslips were then mounted on slides, and digital images were captured using an LSM 700 or LSM 780 ZEISS laser scanning confocal microscope (Carl Zeiss). Image data were processed using the integrated LSM software. Supplementary Table 4 enlists the antibodies used.

### Flow cytometry analysis

To analyze the expression of cell surface TROP2 protein following *NUTM1*-targeted editing of NC cells, cells were detached with cell dissociation buffer (Themo Fisher Scientific, #13151014). The detached cells were then centrifuged and resuspended in FACS Buffer (PBS solution with 1% FBS) t9o a concentration of 2–3 × 10^6^/mL. The cells were incubated with PE anti-human *TACSTD2* (TROP2) antibody (Biolegend, #363804) in PBS containing 1% FBS for 1 h at 4 °C. Thereafter, the cells were washed twice and transferred to a FACS tube (Corning, #352235) for analysis using the BD FACS Lyric.

To analyze the TROP2 binding of sacituzumab following *NUTM1*-targeted editing of NC cells, cells were detached with accutase (Thermo Fisher Scientific, #00-4555-56). The cells were then incubated with 1 μg/mL of control mAb or sacituzumab in PBS containing 1% FBS for 1 h at 4 °C. Thereafter, the cells were washed twice, incubated with a fluorescein isothiocyanate (FITC)-conjugated anti-human secondary antibody (1:100; Jackson ImmunoResearch, #109-095-098), and transferred to a FACS tube for analysis.

### Apoptosis assay

At 72 h after transfection, the proportion of apoptotic cells among the transfected NC cells was assessed using an Annexin V-FITC Apoptosis Detection Kit (Abcam, # ab14085), following the manufacturer’s instructions. Apoptotic cells were analyzed using a BD FACSLyric flow cytometer and the FlowJo software.

### Senescence assay

At 72 h after transfection, the proportion of senescent cells among the transfected NC cells was assessed using the Cell Event Senescence Green Detection Kit (Invitrogen, #C10850), following the manufacturer’s instructions. Senescent cells were visualized under a fluorescence microscope.

### Production of sacituzumab–IFN-**β** mutein

IFN-β muteins, which were purified as described previously ^44^, were produced by Abion Inc. (Seoul, South Korea) using a Chinese hamster ovary cell line ^45^. The genes encoding sacituzumab and sacituzumab–IFN-β mutein were inserted into Freedom pCHO 1.0 expression vectors (Gibco). This Freedom pCHO 1.0 vector was transfected into CHO-S cells (Gibco) using FreeStyle™ MAX reagent (Thermo Fisher Scientific) and OptiPRO™ SFM (Thermo Fisher Scientific). Subsequently, the culture supernatant from CHO-S cells stably expressing sacituzumab and sacituzumab-IFN-β mutein was purified with a MabSelect SuRe™ Protein A resin (Cytiva) using the AKTA avant150 system (Cytiva).

### ADCC assay

The ADCC assay was performed using HCC2429 cells. The cells were seeded in 96-well plates at a density of 2 × 10^4^ cells/well and incubated overnight. The previously established NK-92MI-hCD16a cells stably expressing human CD16a were used as effector cells ^46,47^. HCC2429 cells were incubated with control mAb, IFN-β mutein, sacituzumab, or sacituzumab–IFN-β mutein at a final concentration of up to 100 nM and 2 × 10^4^ effector cells in a CO_2_ incubator for 4 h at 37 °C (E:T ratio = 1:1). HCC2429 cell lysis was measured by detecting the release of lactate dehydrogenase using CytoTox 96® Non-Radioactive Cytotoxicity Assay Kit (DogenBio, # EZ-LDH) following the manufacturer’s instructions. The absorbance of the plates was analyzed using a Spark TM 10M microplate reader (TECAN). For the data analysis, the percentage of specific ADCC was calculated as follows:

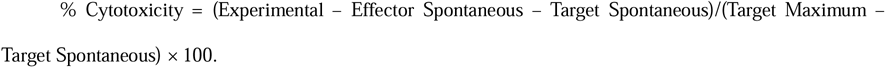

The dose–response curve and EC50 values were estimated using GraphPad Prism 10.

### RNA sequencing

Total RNA was extracted from NC FFPE samples with sufficient remaining tissue using the RNeasy FFPE Kit (Qiagen, #73504). For freshly frozen samples and cell pellets, RNA extraction was performed using a RNeasy Mini Kit (Qiagen, #74004), following the manufacturer’s protocol. The concentration and purity of the extracted RNA were assessed using a Nanodrop® (NanoDrop Technologies Inc.) and a Qubit® 3.0 Fluorometer (Thermo Fisher Scientific, #Q33216). RNA sequencing was performed by Macrogen following the manufacturer’s protocols and reagents. Sample libraries were prepared independently using the TruSeq Stranded mRNA Sample Prep Kit (Illumina, Inc.). The indexed libraries were sequenced on an Illumina NovaSeq platform (Illumina, Inc.) by Macrogen using paired-end (2 × 101 bp) sequencing. The quality of the sequencing data was assessed using FastQC (v0.11.7), and trimming was performed using Trimmomatic (v0.38). High-throughput sequencing data were mapped to the reference genome using HISAT2 (v2.1.0) and Bowtie2 (v2.3.4.1). Following alignment, StringTie (v2.1.3b) was used to assemble the aligned reads into transcripts and estimate their abundance.

### Identification of partner genes for the *NUTM1* fusion gene

RNA was extracted and quantified as previously described. The initial analysis involved qRT-PCR using primer pairs targeting *BRD4* exon 11 and *NUTM1* exon 3. Samples that were not detected by qRT-PCR were subsequently subjected to further analysis using a targeted NGS assay, as described previously^48^. RNA sequencing was performed for samples that were not detected using the aforementioned methods, and fusion partner genes were identified using various fusion detection tools, including Defuse (v0.8.1), FusionCatcher (v1.00), Arriba (v1.2.0), and STAR-Fusion (v1.13.0).

### Transcriptome data analysis

#### Bulk RNA-sequencing data analysis

Gene-level transcription analysis was performed using the DESeq2 (v1.42.0). To investigate the pathway enrichment, hallmark gene sets from the MSigDB database were used. Pathway enrichment analysis was performed using GSVA (v1.50.5), which converts gene expression data into pathway profiles by applying single-sample gene set enrichment analysis (GSEA) within the GSVA framework.

#### Visium data analysis

The files obtained using 10× Visium were analyzed using Seurat (v5.0.1)^49^ in R and Scanpy (v1.10.0rc2)^50^ in Python. Genes were filtered out if they were mitochondrial genes or if the minimum number of spots was ≤3, and cells were filtered out if the minimum number of genes was ≤200. Patients with counts >50,000 were excluded. Subsequently, a normalization procedure was implemented using the SCTransform^51^ method to harmonize the expression profiles across samples and mitigate technical bias. Dimensionality reduction was performed using Principal component analysis followed by UMAP. DEGs were obtained using DESeq2. For GSEA, only significant genes were identified with an adjusted p < 0.05. The rank of the gene list was determined using Wald test statistics.

#### Spot-level annotation and cell-type deconvolution

Cell-type composition per spot was estimated using Conditional Auto Regressive Deconvolution^52^, referencing the “Cell Type Major” and “Cell Type Tumor” categories from the Lung Cancer Atlas Core Atlas^53^. However, initial deconvolution results produced inconsistent classifications between PDT and DT regions depending on the reference category, likely due to the absence of an in-house single-cell reference and the intrinsic spatial resolution limit of Visium (5-10 cells per spot), which results in signal mixing from heterogeneous populations. To address this limitation and improve spatial accuracy, two certified pathologists manually annotated the tumor and normal regions into PDT, DT, PDT-DT mixed, and stromal subtypes including lymphocyte-enriched stroma, bronchiole epithelium, and alveolar septa using the Loupe Browser (v8.0.0), based on histological features. Spots with ambiguous cellular composition (>30% mixed) or low UMI counts were excluded from analysis. These annotations were used for downstream analyses.

#### Cell-of-origin inference

SingleR (v2.8.0)^37^ was used for independent validation of annotated regions via reference-based cell-of-origin classification. Discrepancies between SingleR and manual annotation were reviewed and resolved by pathologist consensus.

### Specimens and histopathological evaluation

The initial 27 patients diagnosed with NC with NUT IHC positivity between 2017 and 2024 at the Samsung Medical Center were selected based on electronic medical records and the results of a previous study.^48^ Further histopathological diagnoses and evaluations were performed by two certified pathologists who reviewed whole slides of FFPE samples. Finally, 24 cases were confirmed as NC after excluding two SMARCA4-deficient tumors and one germ cell tumor with aberrant NUT IHC positivity. Confirmed 24 NC cases were classified into the following subtypes according to their degree of differentiation: UDTs, PDTs, and DTs. PDTs display a poorly differentiated squamous cell carcinoma-like morphology composed of cohesive epithelial tumor cells without keratinization in a solid or lobulated architecture. Tumors with a single focal abrupt keratinization or a squamous pearl were classified as differentiated. Other tumors, such as those with a small round cell morphology without evidence of squamous cell carcinoma features, were classified as UDTs. From the 24 confirmed cases, the available samples were used for qRT-PCR and/or NGS assays to identify *NUTM1* fusion partner genes using both FFPE tissue blocks and fresh frozen samples. This study was reviewed and approved by the Institutional Review Board of Samsung Medical Center (2021-10-097, 2022-11-132, and 2025-03-013).

### Survival analysis

progression-free survival was defined as the time from the start of any treatment administered to the patient until the first documented signs of tumor recurrence or disease progression were confirmed using radiological imaging or during routine follow-up. The R packages survival (v3.7.0), survminer (v0.4.9), and forest (v0.3.0) were used to generate HRs, forest plots, and Cox regression models.

### Statistics and reproducibility

All statistical analyses were performed using R software and GraphPad Prism version 10. Data are presented as mean ± SEM. Statistical significance between the two groups was assessed using an unpaired Student’s two-tailed t-test. Comparisons among three or more groups were performed using one-way ANOVA with Tukey’s multiple comparison test or two-way ANOVA with Tukey’s multiple comparison test, as appropriate. To assess the differences in gene GSVA scores among the UDT, PDT, and DT groups, we used the Kruskal-Wallis test, followed by the Wilcoxon rank-sum test with Benjamini-Hochberg correction for multiple comparisons. Statistical significance was set at p < 0.05. Statistical significance is annotated directly on the plots using asterisks (*p < 0.05, **p < 0.01, ***p < 0.001, and ****p < 0.0001). The specific statistical tests applied in each experiment are indicated in the respective figure legends.

### BioRender

Figures were created with BioRender.com

## Data availability

## Acknowledgments

We thank Christopher Alexander French (Harvard Medical School) for providing–10-15 and JCM cells. This work was supported by the National Research Foundation of Korea (NRF) grants funded by the Korean government (Ministry of Science and ICT) (RS-2022-NR070196, RS-2023-00217189, RS-2024-00347894) and by a grant from the National R&D Program for Cancer Control, Ministry of Health & Welfare, Republic of Korea (RS-2023-CC138390).

## Author contributions

Y.L.C. and M.S.L. conceived the original idea and supervised the study. J.Y.C. and D.S.K. performed primary experiments. J.H.O. performed spatial transcriptomic data analysis. H.G.P. performed ADCC assay. H.L.Y. provided Cas9-eVLPs. S.L. provided sacituzumab–IFN-β mutein. Y.L. and Y.L.C. performed pathological examination. M.J.S. collected the clinical samples. H.Y.J., J.Y.S., E.S.C., K.S.J. analyzed and interpreted the data. D.S.K., Y.K.S., S.H.L. and C.F provided scientific guidance for the research and critically reviewed the manuscript. J.Y.C. and M.S.L. wrote and revised the manuscript. The order of co-first authors was determined based on the complexity and extent of the experiments conducted. All authors reviewed and approved the manuscript.

## Additional information

**Supplementary information** The online version contains supplementary material available at ∼

