## Supplementary Material for "CRISPR-Induced NUT Suppression Promotes Differentiation and Enhances Trop2-Targeted Immunocytokine Response in NUT Carcinoma"

**­****Supplementary Information**

Supplementary Information includes:

- Supplementary Figures. 1-8
- Supplementary Tables. 1-4

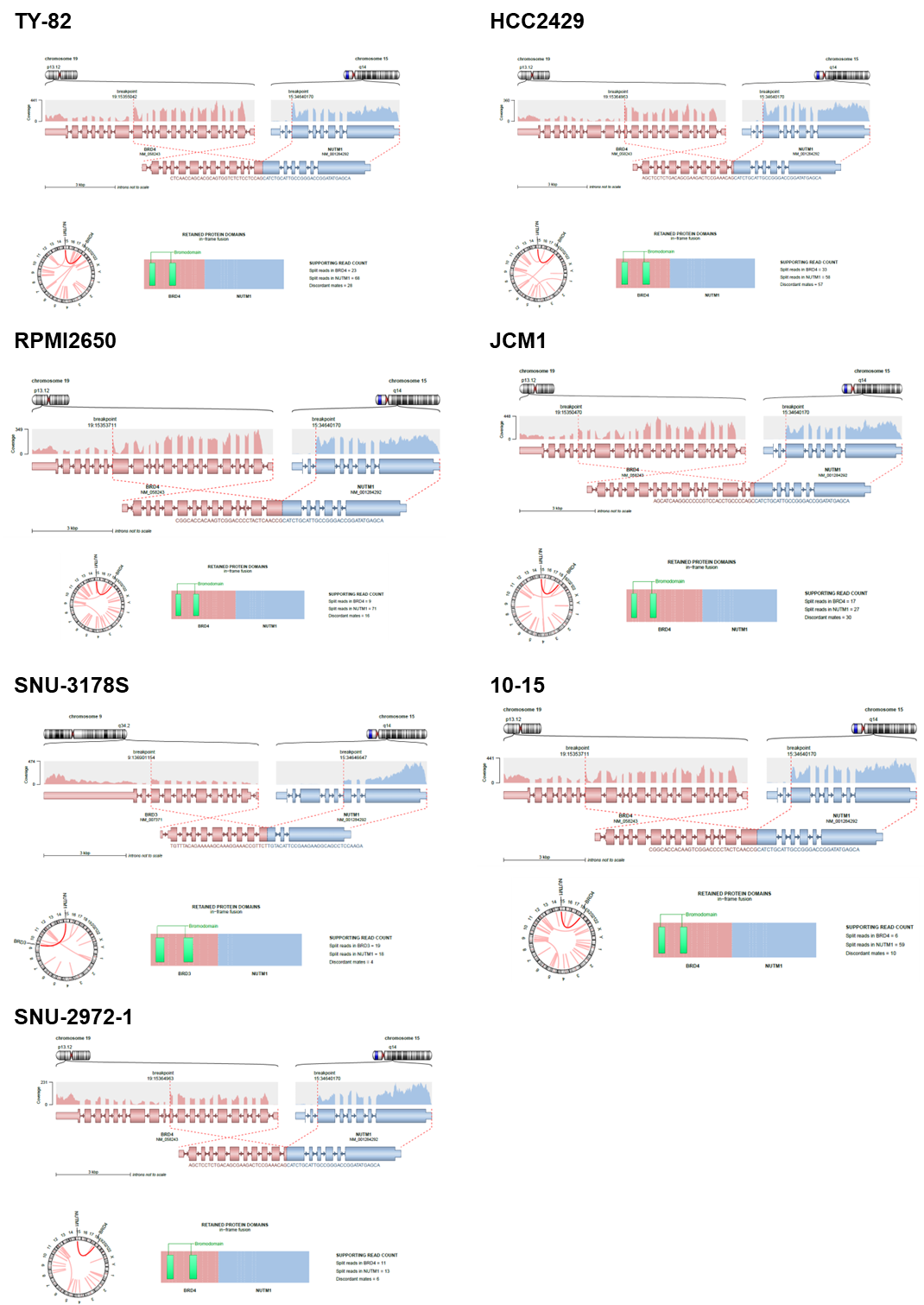

**Supplementary Figure 1. Detection of Fusion Partners in NC Cell Lines Using RNA Sequencing.**
Figure presenting fusion gene detection results from RNA sequencing analysis of NC cell lines using the Arriba package. Each panel highlights fusion events, displaying the involved genes, fusion breakpoints, and supporting read counts. Key elements include the genomic loci of fusion partners, RNA-seq read coverage, and chromosomal translocation points. These results reveal critical fusion events involving *NUTM1*, contributing to the molecular characterization of NUT carcinoma.

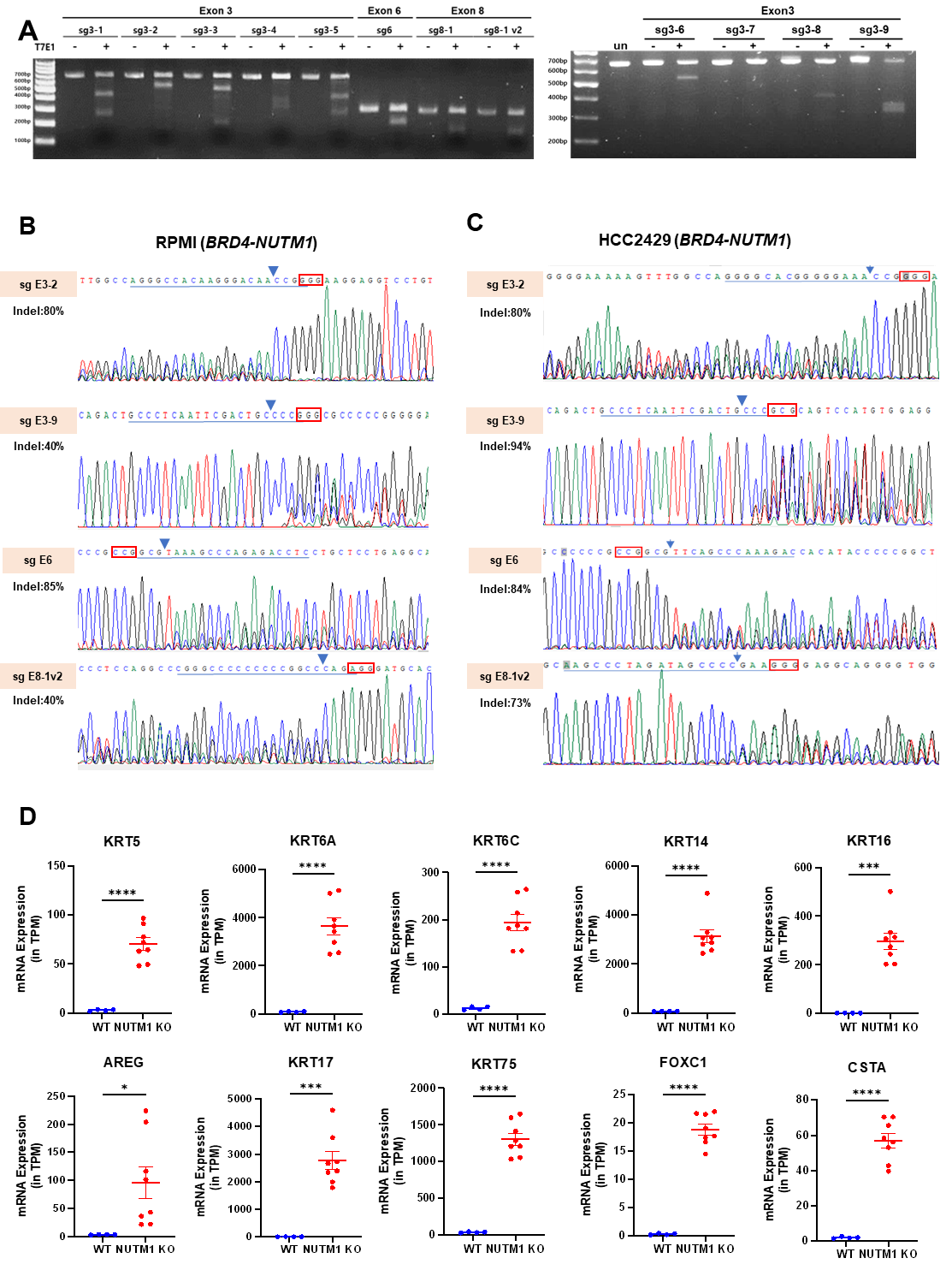

**Supplementary Figure 2. Validation of sgRNA Efficiency and NUTM1 Editing promotes differentiation.**

(A) T7 endonuclease 1 assay results demonstrating sgRNA efficiency across different sgRNAs targeting NUTM1. Gel electrophoresis images show cleavage patterns indicative of sgRNA activity. (B,C) Sanger sequencing validating indel formation at the target sites. Red boxes indicate PAM sequences and blue arrow indicates the regions with sequence alterations. Sequencing was performed for sgRNAs E3-2, E3-9, E6, and E8-1v2 on RPMI and HCC2429 cell lines. (D) Dot plot depicting the increased mRNA expression of differentiation-related genes in NUTM1-edited cells, suggesting a shift towards a more differentiated, less oncogenic phenotype. Data are presented as the mean ± SEM. Statistical significance between the 2 groups was assessed using an unpaired Student’s 2-tailed t test. *p < 0.05, ***p < 0.001, ****p < 0.0001.

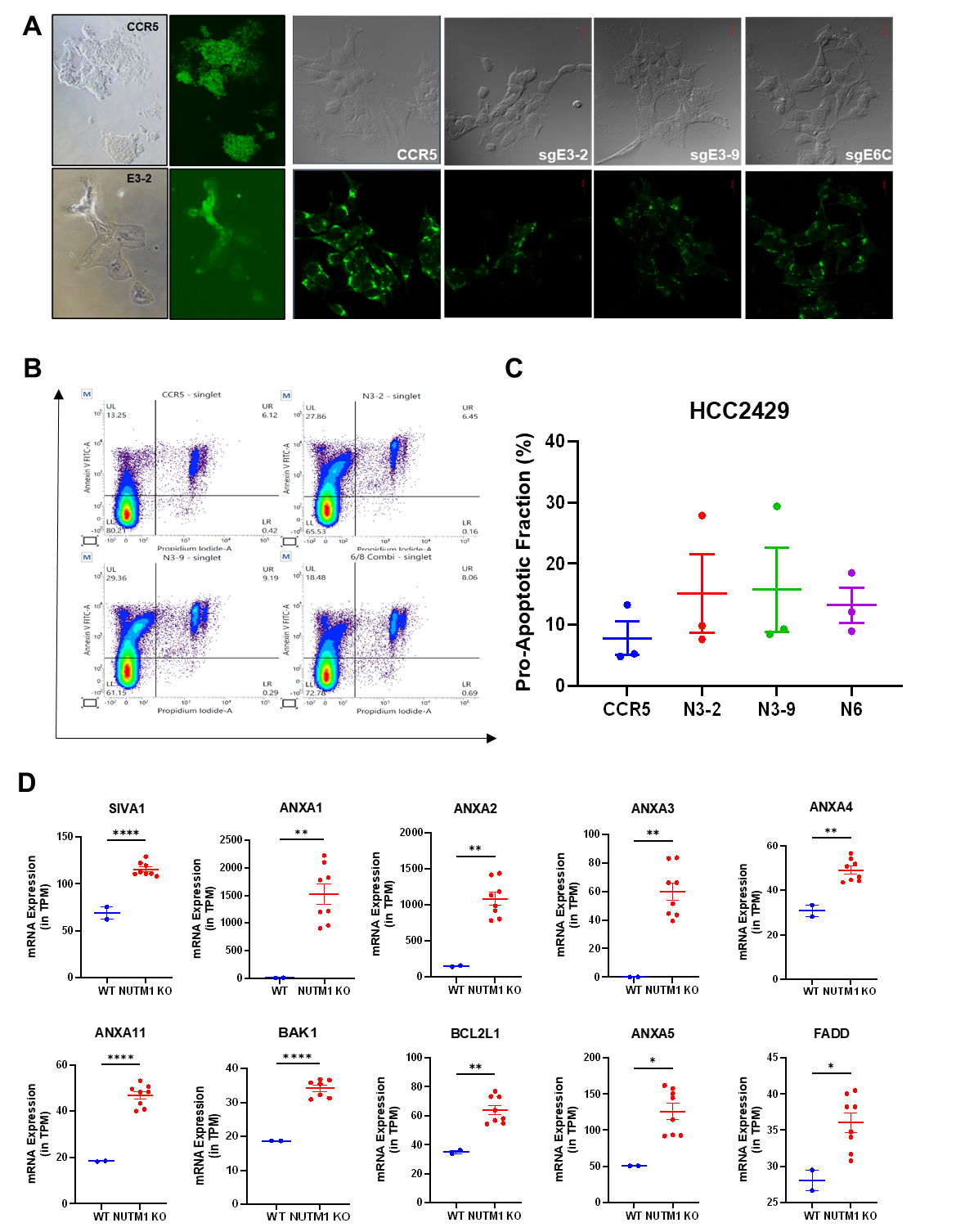

**Supplementary Figure 3. Senescence and Apoptosis Assays Indicate that Morphological Changes in NUTM1-Edited Cells Result from Differentiation.**

(A) Representative images of NUTM1-edited cells (sgE3-2, sgE3-9, and sgE6) and control CCR5-edited cells under a fluorescence microscope. Brightfield and GFP fluorescence images show no significant increase in GFP-positive cells, indicating the absence of senescence. (B) Flow cytometry analysis of apoptosis in NUTM1-edited and control cells. Annexin V and Propidium Iodide staining were used to assess apoptotic populations. The graph shows no significant increase in overall apoptosis but a slight rise in the pro-apoptotic fraction in edited cells. (C) Dot plot depicts pro-apoptotic fractions across different sgRNAs targeting NUTM1. Data are presented as the mean ± SEM. No statistical significance observed. (D) Dot plot showing a slight increase in the mRNA expression of pro-apoptotic genes in NUTM1-edited cells. Data are presented as the mean ± SEM. Statistical significance between the 2 groups was assessed using an unpaired Student’s 2-tailed t test. *p < 0.05, **p < 0.01, ****p < 0.0001.

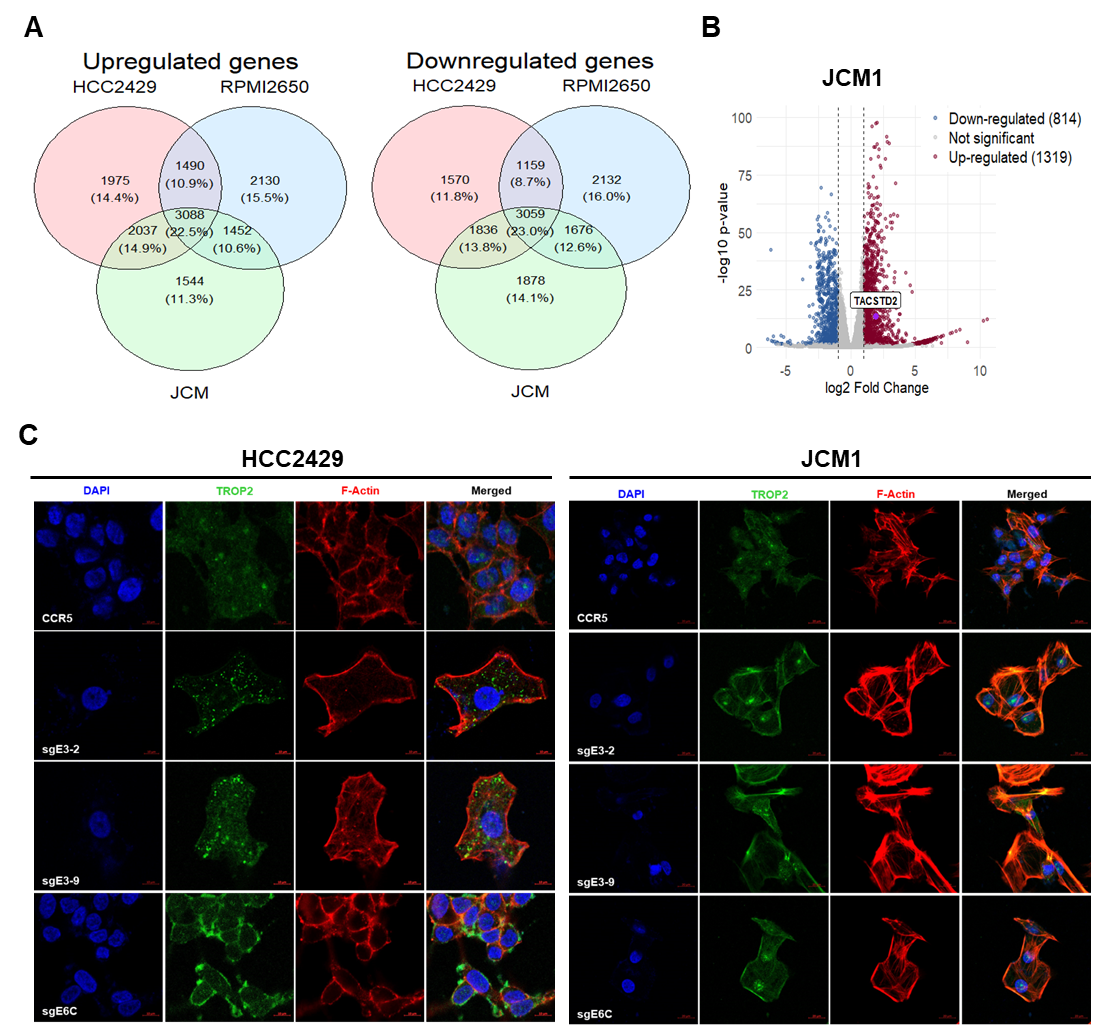

**Supplementary Figure 4. *TACSTD2* expression increases in NUTM1-editied NC cell lines**

(A) Venn diagrams showing the number of overlapping upregulated and downregulated genes in the NUTM1-edited NC cell line. (B) Volcano plot depicting differentially expressed genes between CCR5 and NUTM1-edited JCM cells. The TACSTD2 gene is highlighted as significantly upregulated. (C) Immunofluorescence analysis of cytoskeletal changes in CCR5 and NUTM1-edited cells. Cells were stained for DAPI (nuclei, blue), TROP2 (green), and F-actin (red). NUTM1-edited cells exhibited enhanced stress fiber formation, indicating cytoskeletal reorganization associated with cellular differentiation. Scale bars represent 5 µm.

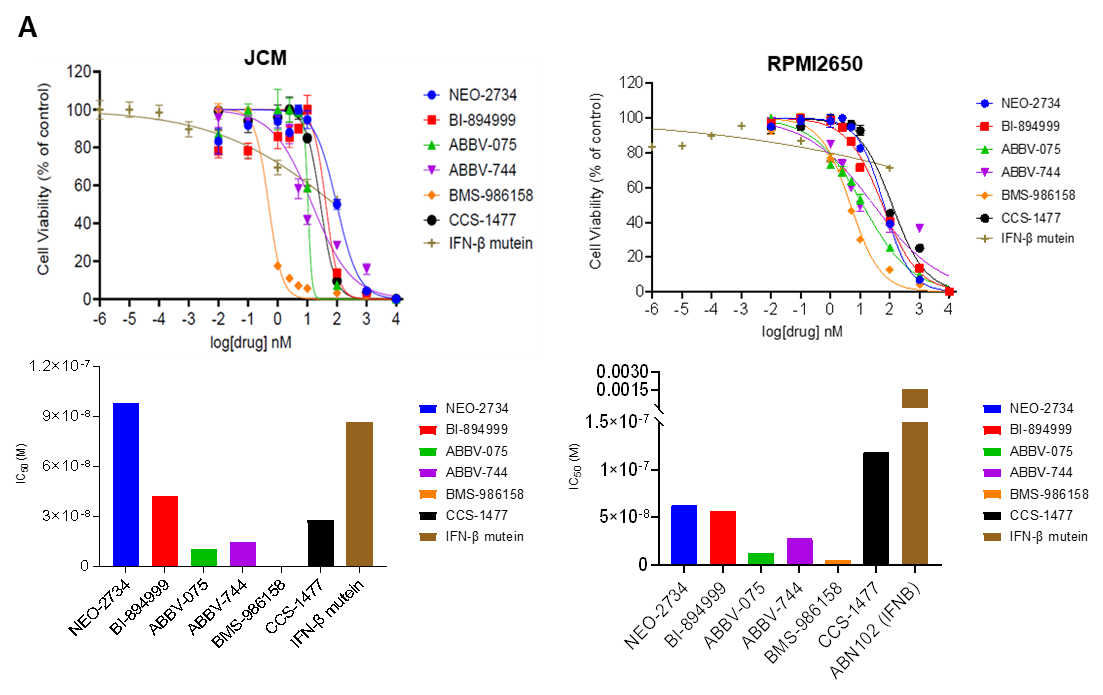

**Supplementary Figure 5. Evaluation of BET Inhibitors and IFN-β Mutein in Various NC Cell Lines**

(A) Dose-response curves showing cell viability following treatment with various BET inhibitors (NEO-2734, BI-894999, ABBV-075, ABBV-744, BMS-986158, CCS-1477) and IFN-β mutein in NC cell lines. Drug efficacy was assessed by plotting cell viability (%) against the log of drug concentration (nM).

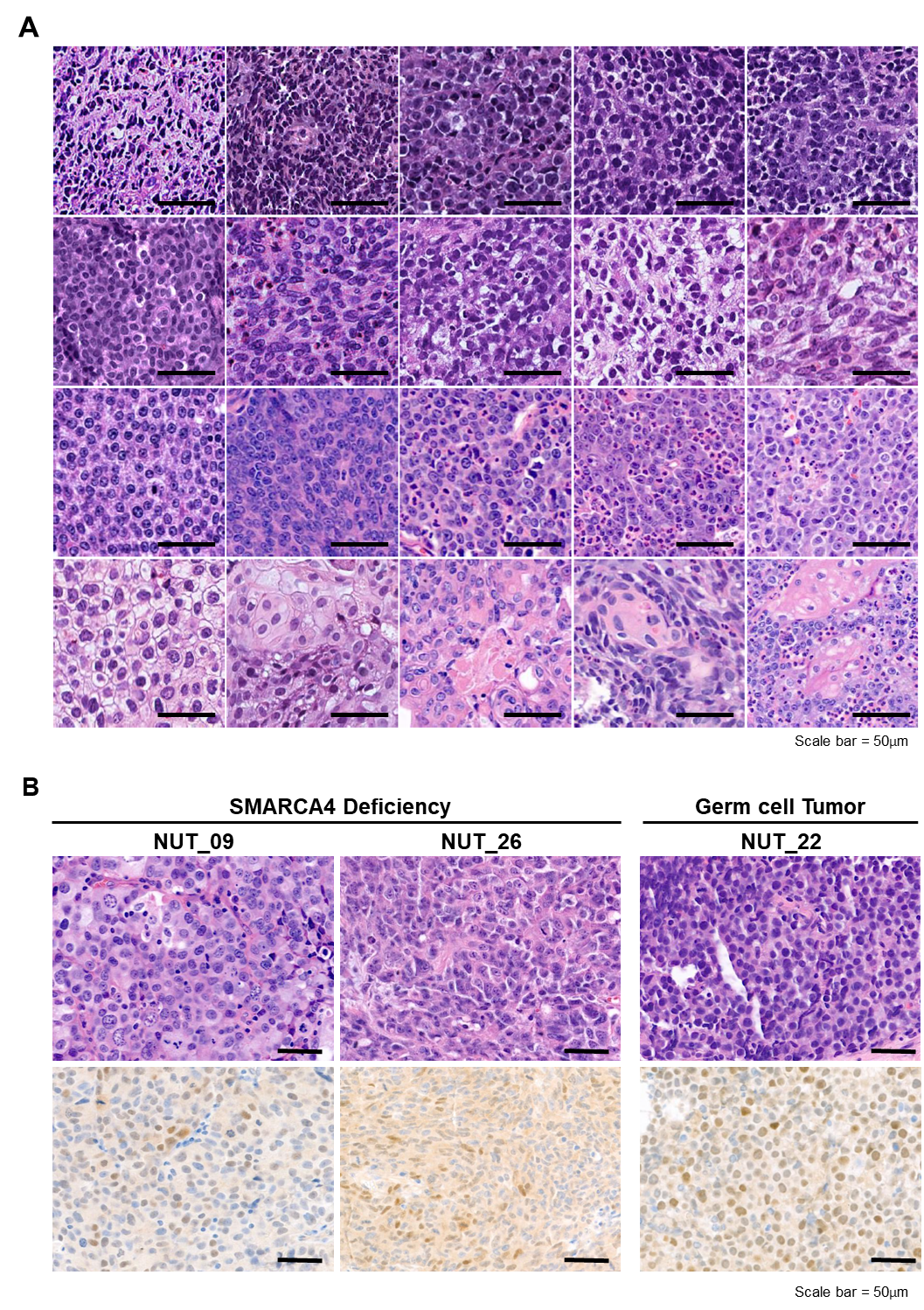

**Supplementary Figure 6. Histological Classification of NC Patients and Diagnostic Confirmation through Histopathological Evaluation**

(A) H&E images of tumors from 20 NC patients were evaluated for histopathological characteristics, showing varying degrees of differentiation. Scale bars represent 50 µm. (B) H&E and NUT IHC images of three atypical NC cases are shown. These cases exhibited unusual patterns, including weak cytoplasmic staining or the absence of nuclear speckled expression. Two cases demonstrated BRG loss, indicating SMARCA4-deficient tumors, while the third case was diagnosed as a mediastinal germ cell tumor. These cases were excluded from the cohort.

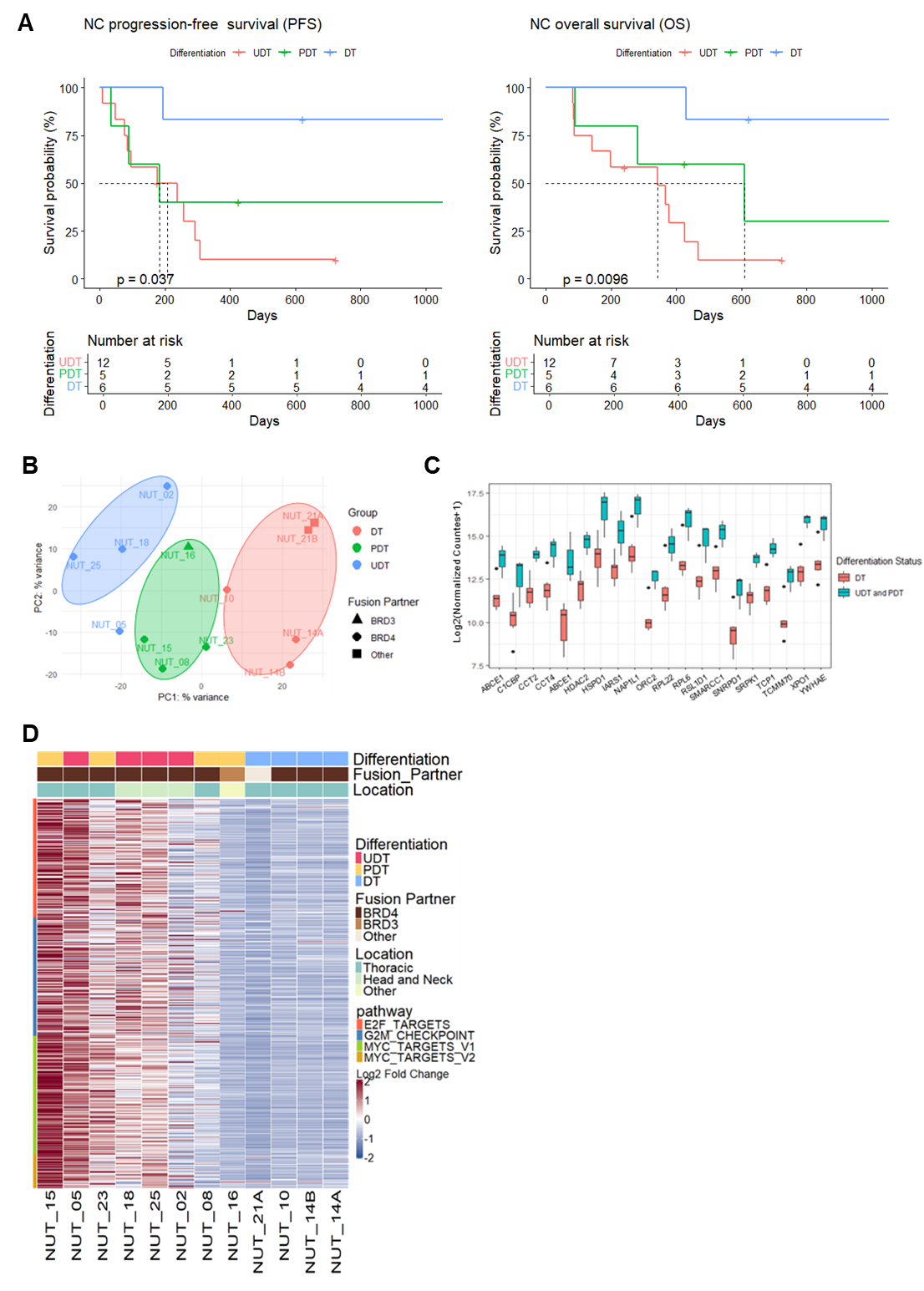

**Supplementary Figure 7. Prognostic Impact of Tumor Differentiation on Survival and Pathway Enrichment**

(A) Kaplan-Meier curves of progression-free survival (PFS) and overall survival (OS) for patients with DT, PDT, and UDT. Patients with DTs showed significantly better PFS (p = 0.037) and OS (p = 0.0096) compared to those with PDTs and UDTs. (B) Principal component analysis PCA plots of transcriptomics data of patients with NC. (C) Expression of Hallmark MYC pathway genes stratified as differentiation status. (D) Heatmap showing pathway enrichment analysis, focusing on proliferative pathways associated with NC, including the G2M checkpoint, E2F targets, and MYC targets. The heatmap also highlights tumor differentiation status, fusion partners (BRD4, BRD3, or others), and tumor location (thoracic, head and neck, or other). PDTs and UDTs demonstrated upregulation of proliferative pathways, reflecting aggressive tumor phenotypes.

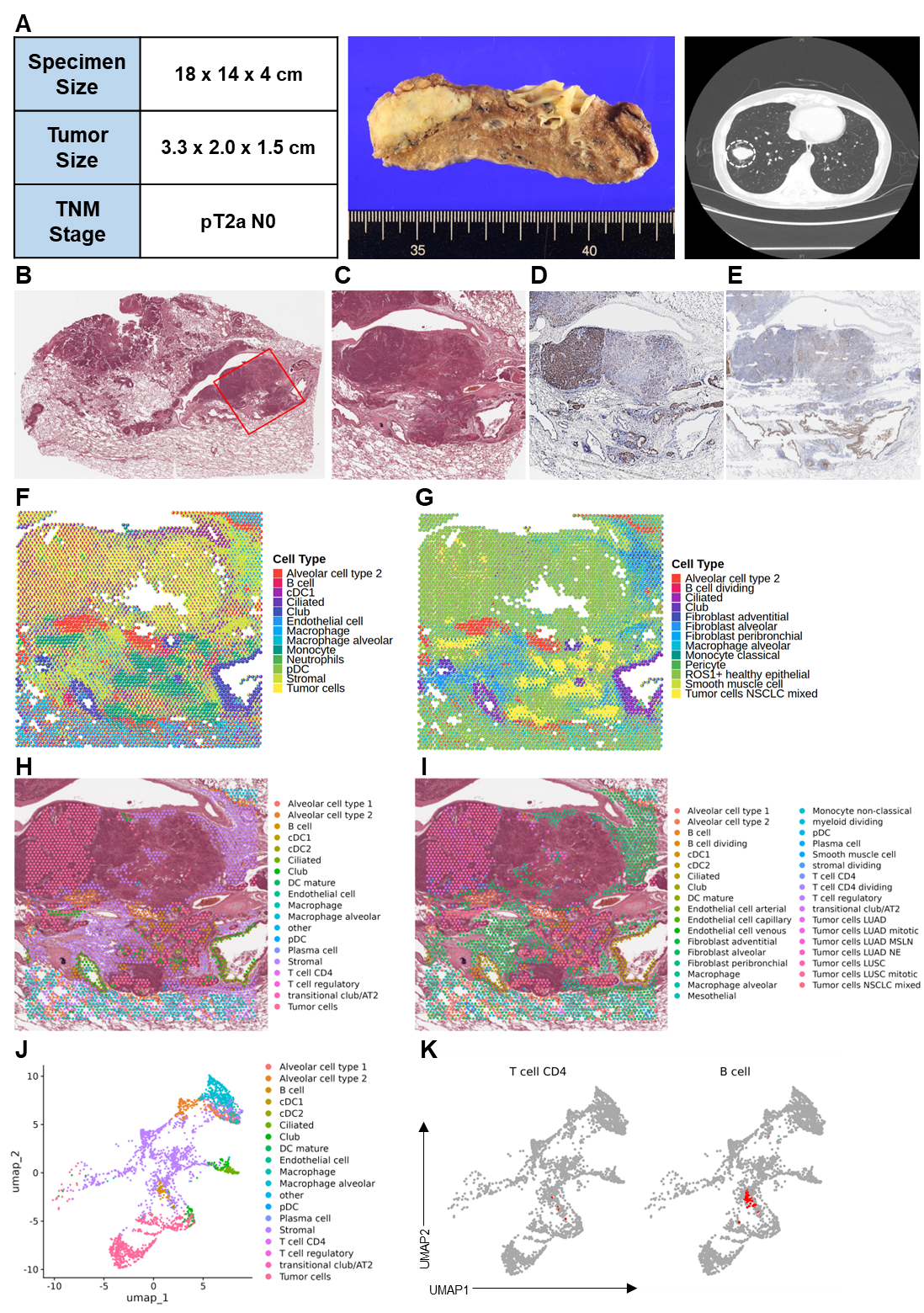

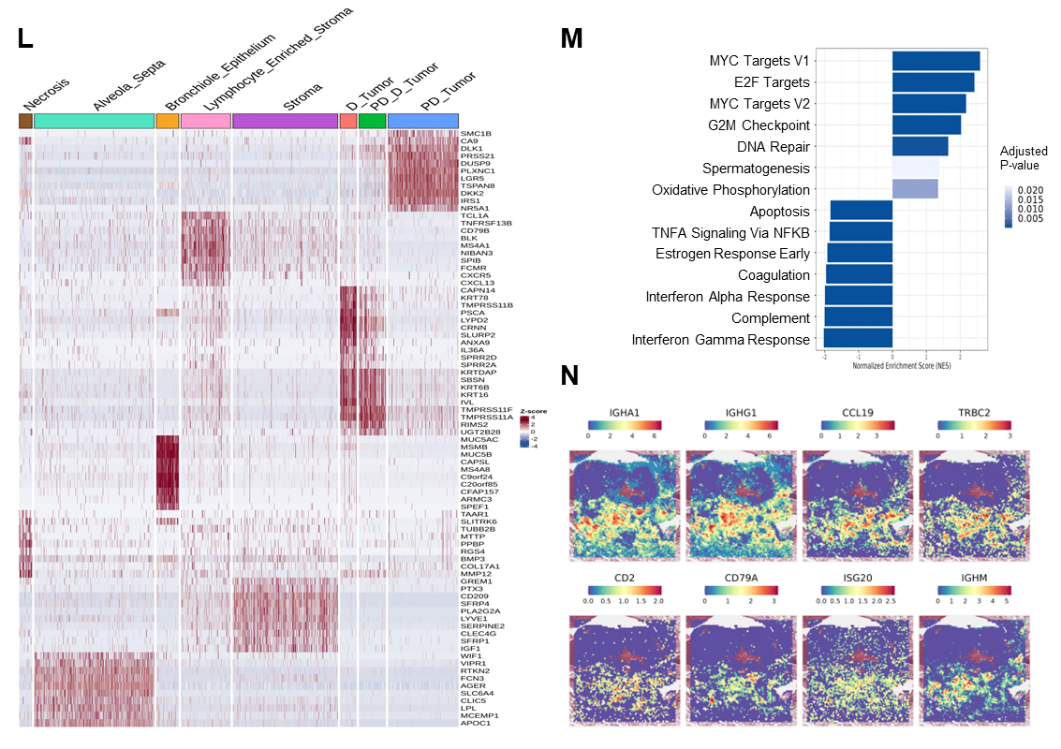

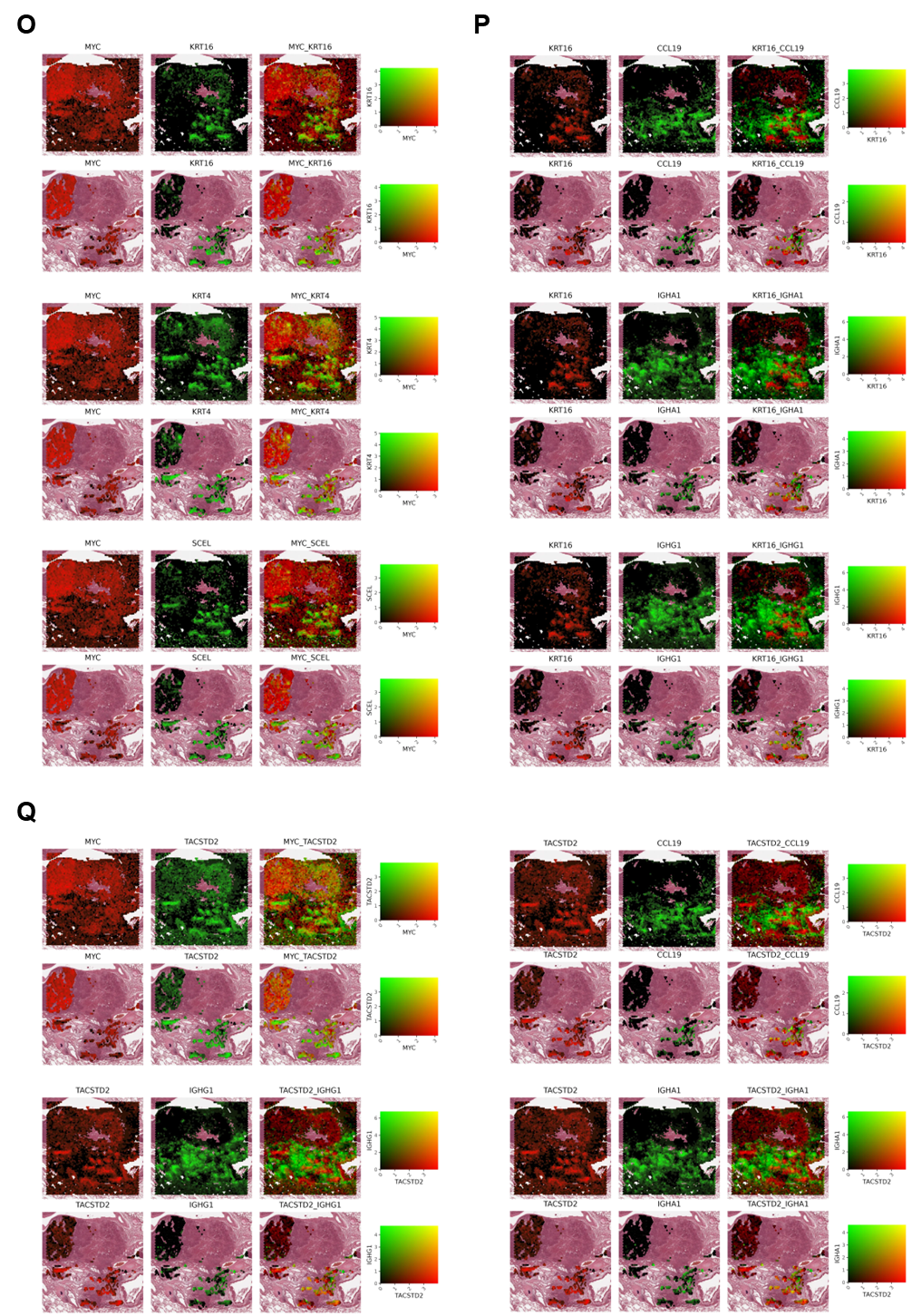

**Supplementary Figure 8. Integrated spatial analysis of differentiation, proliferation, and immune activity in NC Patient.**

(A) Specimen information for NUT14 Patient, including gross specimen, CT scan images showing tumor location, and details on specimen size, tumor size, and TNM stage. (B) Low-magnification H&E image highlighting the region of interest (ROI) for spatial transcriptomics (indicated by the red box). (C) ROI used in the actual experiment as observed in Loupe Browser. (D, E) NUT and TROP2 IHC staining of sample. (F, G) Cell-type deconvolution was performed using CARD on Visium spatial transcriptomic data. The Single-cell Lung Cancer Atlas (LuCA) dataset served as the reference, utilizing the 'Cell Type Major' and 'Cell Type Tumor' annotations from the Core Atlas, respectively. (H, I) Cell-of-origin analysis was conducted using the SingleR package, based on the same LuCA reference dataset, applying the 'Cell Type Major' and 'Cell Type Tumor' categories, respectively. (J) UMAP visualization of the 'Cell Type Major' annotation. (K) UMAP shows that immune-enriched clusters, including CD4+ T cells and B cells, are located near DT regions. (L) Heatmap illustrating distinct gene expression profiles across annotated regions, demonstrating spatial and molecular heterogeneity within the NC sample. (M) GSEA analysis showed that PDT and PDT-DT were enriched for proliferative pathways, whereas DT was enriched for immune-related pathways. (N) Spatial feature plots showing the localization of immune markers (IGHA1, CCL19, IGHG1, TRBC2 and etc.) in DT regions, indicating a more immune-active microenvironment compared to PDT regions. (O) Co-localization of proliferation marker (MYC) with differentiation markers (KRT4, KRT16, and SCEL). MYC was enriched in PDT regions, while differentiation markers were predominant in DT regions, indicating inverse expression patterns and reduced proliferation in DTs. (P) Co-localization of KRT16 with immune markers (CCL19, IGHA1, IGHG1) in DT regions, suggesting an association between differentiation and immune activity. (Q) Co-localization of TACSTD2 with immune markers in DT regions, supporting the presence of immune-enriched microenvironments that may enhance responsiveness to TROP2–IFN-β mutein immunocytokine therapy.

**Supplementary Tables**

| **Cell line** | **Patient diagnosis** | **NC translocation** |
| --- | --- | --- |
| TY82 | 22-year female; thymus, NUT Midline Carcinoma | t(15;19)(q14;p13); BRD4::NUT (ex 13::ex 3)  [NM 001284292] |
| RPMI2650 | 52-year male; nose (nasal septum),  NUT Midline Carcinoma | t(15;19)(q14;p13); BRD4::NUT (ex 14::ex 3) |
| HCC2429 | 34-year female; lung, NUT Midline Carcinoma | t(15;19)(q14;p13); BRD4::NUT (ex 11::ex 3) |
| JCM | NUT Midline Carcinoma | t(15;19)(q14;p13); BRD4::NUT (ex 16::ex 3) |
| 10-15 | NUT Midline Carcinoma | t(15;19)(q14;p13); BRD4::NUT (ex 14::ex 3) |
| SNU-3178S | 33-year ever-smoking female; lung with multiple bone metastases (pleural effusion),  NUT Midline Carcinoma | t(15;9)(q14;p34.2A); BRD3::NUT ( ex 10::ex 6) (Annals of Oncology, 2017) |
| SNU-2972-1 | 34-year male, sinonasal and intrathoracic with pleural effusion  NUT Midline Carcinoma | t(15;19)(q14;p13); BRD4:: NUT (ex 11::ex 3) |
| PER-403 | 11-year female; thymus, poorly differentiated intrathoracic squamous carcinoma | t(15;19)(q14;p13.1): BRD4::NUT (ex 11::ex 3) (Kees et al, 1991) |
| PER-624 | 16-year female; lung, poorly differentiated aggressive lung carcinoma, complex | t(6;19)(q13;p13.1): Cryptic BRD4::NUT (ex 15::ex 3) (Thompson-Wicking et al, 2013) |
| PER-704 | 8-year male; larynx, poorly differentiated laryngeal carcinoma | t(15;19)(q14;p13.1): BRD4::NUT (ex 15:: ex 3) |

**Supplementary Table 1. Lists of cell lines*.***

Each cell line's patient origin and fusion partner are described in detail.

| **Target Exon** | **sgRNA ID** | **sgRNA sequence** | **PAM Sequence** |
| --- | --- | --- | --- |
| 3 | **sg E3-1** | TCAAAGTCAAGACAGAAGGG | GGG |
| 3 | **sg E3-2** | AGGGCCACATGGGACAACCG | GGG |
| 3 | **sg E3-3** | CCAGCTCACTGTTGGTGACA | GGG |
| 3 | **sg E3-4** | (-) GCCCGGGGCAGTCGAATTGA | GGG |
| 3 | **sg E3-5** | ATTCTGCCCTCTAAGGCTGT | TGG |
| 3 | **sg E3-6** | (-) GAGAGCATCAGAGGGTTGTC | TGG |
| 3 | **sg E3-7** | GATGGGGGCCCTTGCCTCAG | TGG |
| 3 | **sg E3-8** | (-) GATAAAGTTCTGAGTTTGAG | AGG |
| 3 | **sg E3-9** | GCCCTCAATTCGACTGCCCC | GGG |
| 6 | **sg E6** | (-) TCTGGGCTTTACGCTGACGC | CGG |
| 8 | **sg E8-1** | CAAGCCCTAGATAGCCCCAG | AGG |
| 8 | **sg E8-2v2** | AAGCCCTAGATAGCCCCAGA | GGG |
| *E6C - 1:1 mixture of sg E6 and sg E8-1v2 | | | |

**Supplementary Table 2. Lists of sgRNA sequences targeting *NUTM1.***

Table of *NUTM1*-targeting sgRNAs, detailing target exons, sgRNA IDs, sequences, and PAM sequences. The group referred to as E6C represents the 1:1 mixture of sgE6 and sgE8-1v2. Antisense sgRNA sequences are indicated by (-).

| **Primer name** | **Sequence (5'->3')** |
| --- | --- |
| rt_TACSTD2_F | CAAGTGTCTGCTGCTCAAGG |
| rt_TACSTD2_R | TAGAGGCCATCGTTGTCCAC |
| rt_NUTM1 E3_F | GTCAGTGGCAGCGTTACAAA |
| rt_NUTM1 E3_R | CGAAGCACTGGGATAAGAAAAC |
| rt_GUSB_F | CTCATTTGGAATTTTGCCGATT |
| rt_GUSB_R | CCGAGTGAAGATCCCCTTTTTA |
| CCR5_Indel_F | CCCAGTGGGACTTTGGAAATA |
| CCR5_Indel_R | ACCAGCCCCAAGATGACTATC |
| NUTM1_E3_Del_F | ACCGGATATGAGCATGAAACCT |
| NUTM1_E3_Del_R | TGTGTCAGGACTCTGGGATAGG |
| NUTM1_E6_Del_F | GATTGCTGAGCTTACAGAGCTGG |
| NUTM1_E6_Del_R | CACCAGCCATTCCATGATGTC |
| NUTM1_E8-1_Del_F | TCCTTCTGGTTCTGTTGAGGATG |
| NUTM1_E8-1_Del_R | TATAACCTTCCCAGACCCTCTCC |

**Supplementary Table 3. Primer sequences for qRT-PCR and indel analysis.**

| **Antibodies** | **Host** | **Dilutions** | **Manufacturer** | **Cat no.** |
| --- | --- | --- | --- | --- |
| *Primary antibodies* |  |  |  |  |
| NUT (C52B1) | Rabbit | 1:500 (WB) 1:100 (IF) | Cell signaling Technology | #3625 |
| c-Myc | Rabbit | 1:1000 (WB) | Cell signaling Technology | #9402 |
| p63 (N2C1) | Rabbit | 1:2000 (WB) | GeneTex | #GTX102425 |
| SOX2 | Rabbit | 1:4000 (WB) | GeneTex | #GTX101506 |
| β-Actin (C4) | Mouse | 1:1000 (WB) | Santa Cruz Biotechnology | #sc-47778 |
| TACSTD2 | Rabbit | 1:1000 (WB)  1:100 (IF) | Sigma-Aldrich | #HPA055067 |
| *Secondary antibodies* |  |  |  |  |
| Goat Anti-Rabbit IgG H&L (HRP) | Goat | 1:5000 (WB) | Abcam | #Ab6721 |
| Goat Anti-Mouse IgG H&L (HRP) | Goat | 1:5000 (WB) | Abcam | #Ab6789 |
| Goat anti-Rabbit IgG (H+L) Cross-Adsorbed  Secondary Antibody, Alexa Fluor™ 568 | Goat | 1:1000 (IF) | Invitrogen | #A-11011 |
| F(ab')2-Goat anti-Rabbit IgG (H+L) Cross-Adsorbed   Secondary Antibody, Alexa Fluor™ 488 | Goat | 1:1000 (IF) | Invitrogen | #A-11070 |

**Supplementary Table 4. Antibodies used for immunoblotting and immunofluorescence assays.**
